# Massively parallel single-cell B-cell receptor sequencing enables rapid discovery of diverse antigen-reactive antibodies

**DOI:** 10.1101/660472

**Authors:** Leonard D Goldstein, Ying-Jiun J Chen, Jia Wu, Subhra Chaudhuri, Yi-Chun Hsiao, Kellen Schneider, Kam Hon Hoi, Zhonghua Lin, Steve Guerrero, Bijay S Jaiswal, Jeremy Stinson, Aju Antony, Kanika Bajaj Pahuja, Dhaya Seshasayee, Zora Modrusan, Isidro Hötzel, Somasekar Seshagiri

## Abstract

Using high-throughput single-cell B-cell receptor sequencing (scBCR-seq) we obtained accurately paired full-length heavy- and light-chain variable domains from thousands of individual B cells in a massively parallel fashion. We sequenced more than 250,000 B cells from rat, mouse and human repertoires to characterize their lineages and expansion. In addition, we immunized rats with chicken ovalbumin and profiled antigen-reactive B cells from lymph nodes of immunized animals. The scBCR-seq data recovered 81% (n = 56/69) of B-cell lineages identified from hybridomas generated from the same set of B cells that were subjected to scBCR-seq. Importantly, scBCR-seq identified an additional 710 candidate lineages that were not recovered as hybridomas. We synthesized, expressed and tested 93 clones from the identified lineages and found that 99% (n = 92/93) of the clones were antigen-reactive. Our results establish scBCR-seq as a powerful tool for antibody discovery.

## INTRODUCTION

Antibody diversity is an important feature of the adaptive immune system. B cells produce a diverse array of antibodies by rearranging variable (V), diversity (D) and joining (J) immunoglobulin (Ig) germline gene segments^1–3^. Somatic hypermutation and class switching add to antibody diversity. A mature antibody consists of two identical heavy chains linked through disulphide bonds and two identical light chains each linked to one of the heavy chains, generating two identical antigen binding sites formed by the first immunoglobulin domain of each chain combined^2^. The heavy and light chains are encoded in separate gene loci and each B cell normally expresses a single functional heavy and light chain sequence.

Next-generation sequencing (NGS) has been applied to understand the diversity of the variable regions of heavy (VH) and light chains (VL) that determine the antigen specificity of the antibodies. Until recently, the majority of high-throughput sequencing approaches produced unpaired VH and VL repertoires, as generating paired information requires obtaining data at the individual cell level^4^. Recently, techniques using fabricated microfluidic devices or axisymmetric flow-focusing devices and water-in-oil emulsions have demonstrated the potential for obtaining VH-VL pairing information in a high-throughput manner^5–7^. However, limitations in sequencing technologies allow only partial variable region sequences to be obtained using these methods, and clonal size information is partially lost. High-throughput techniques that yield full-length variable region sequence information from individual B cells would enable routine application of large-scale immune repertoire sequencing to antibody discovery and detailed repertoire characterization.

Here we describe the application of high-throughput single-cell sequencing to obtain the VH and VL sequences for antibodies from individual human, rat and mouse B cells. We developed a bioinformatics framework to analyze the sequence data and identify accurate VH and VL pairing. Further, we show the utility of the technique for antibody discovery by expressing and testing predicted antigen-reactive antibody sequences. We demonstrate the potential of direct sequencing of individual antigen-reactive B cells to rapidly generate a large and diverse panel of antigen-specific antibody variable regions and thus expand immune repertoire sampling and expedite antibody discovery processes.

## RESULTS

### Massively parallel sequencing of B-cell receptors from individual cells

We analyzed >250,000 individual IgG^pos^ B cells from three human donors and two mice, and IgM^neg^ B cells from two rats using emulsion-based encapsulation, cDNA generation and sequencing. Briefly, we generated 5’ barcoded cDNA from thousands of individual B cells in parallel, and amplified the VH and VL regions using custom primers while retaining the cell barcode (Fig. 1, Supplementary Fig. 1-2, and Methods). The 5’ barcoded VH and VL domain-encoding cDNAs were sheared and converted into sequencing-ready libraries by addition of appropriate adapter oligonucleotides (Supplementary Fig. 1). The library construction method involves 3’ cDNA shearing after amplification to create a set of fragments with variable 3’ end, while retaining the 5’ end for all fragments. This resulted in sequencing reads with constant 5’ sequence and variable 3’ sequence, allowing *de novo* assembly of full-length VH and VL sequences from short-read data (2 x 150 bp). We devised a computational pipeline for cell detection, *de novo* contig assembly, variable domain annotation, and pairing of full-length VH and VL sequences (Fig. 2a, Supplementary Fig. 3). Assembled VH and VL sequences were parsed for framework and complementarity-determining regions (CDR) and filtered for open reading frames encoding the entire variable regions. As expected, read coverage was highest at the constant 5’ end and more variable at the 3’ end (Fig. 2b, Supplementary Fig. 4). For this reason, we only considered the first 4 codons of the framework 4 region, ensuring that a minimum of 4 codons match germline sequences without frameshifts immediately following the third CDR in both chains (CDR-H3 and L3), providing enough sequence information to determine CDR-H3 and L3 boundaries with high confidence. The median number of reads supporting assembled contigs was at least 175, 213, and 55 throughout the CDR-L3 region, and at least 3, 6, and 11 throughout the CDR-H3 region for rat, mouse and human B-cell repertoires, respectively (Fig. 2b, Supplementary Fig. 4). Cells with complete VH and VL domains were filtered by requiring a minimum number of reads (10 and 100 for VH and VL, respectively) and requiring one dominant VH and VL contig (≥ 80% read support; Fig. 2c, Supplementary Fig. 5). Pairing efficiency, defined as the percentage of cells with at least one detected VH and VL, was 48–94%, depending on sample type, and 48–74% of cells with pairing information passed quality filters (Fig. 2d). Variation in pairing and filtering efficiency may be due, in part, to differences in heavy- and light-chain transcript levels, primer efficiency and primer coverage. Overall, a total of 261,539 sequenced B cells yielded high-quality VH-VL pairing information for 116,115 cells from rat (n = 30,380), mouse (n = 9,459) and human (n = 76,276) B-cell repertoires (Fig. 2d, Supplementary Fig. 3, Supplementary Tables 1–11).

**Figure 1.**
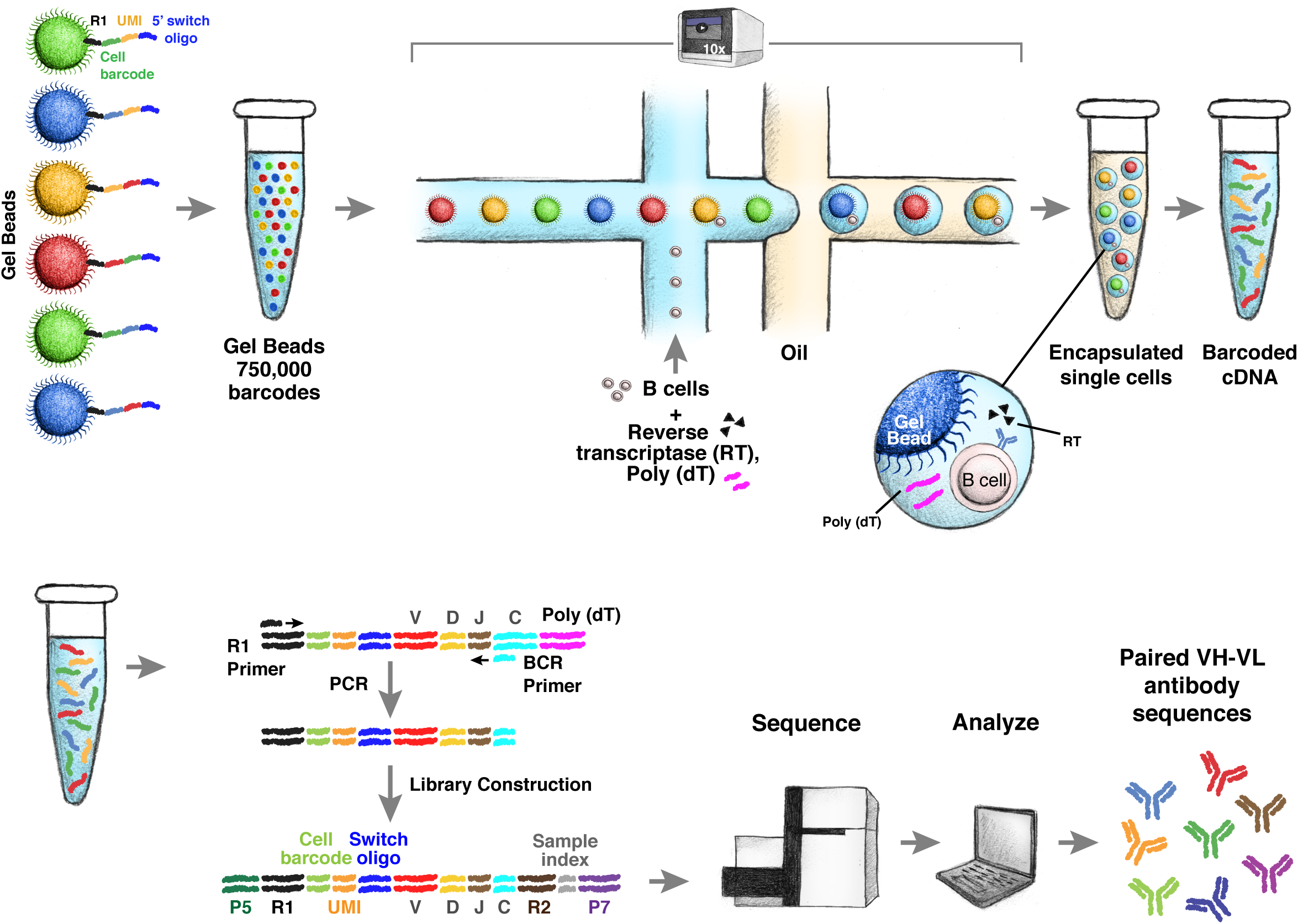
Schematic of single B cell capture, library construction and sequencing. Chromium controller (10x Genomics, Pleasanton, CA) was used to capture single cells along with gel beads containing the 5’ switch oligo with cell barcode, reverse transcriptase and poly(dT) primer. The barcoded cDNAs were then converted into a library and sequenced (Illumina, San Diego, CA). Heavy- and light-chain sequences for individual cells were analyzed using a custom computational pipeline.

**Figure 2.**
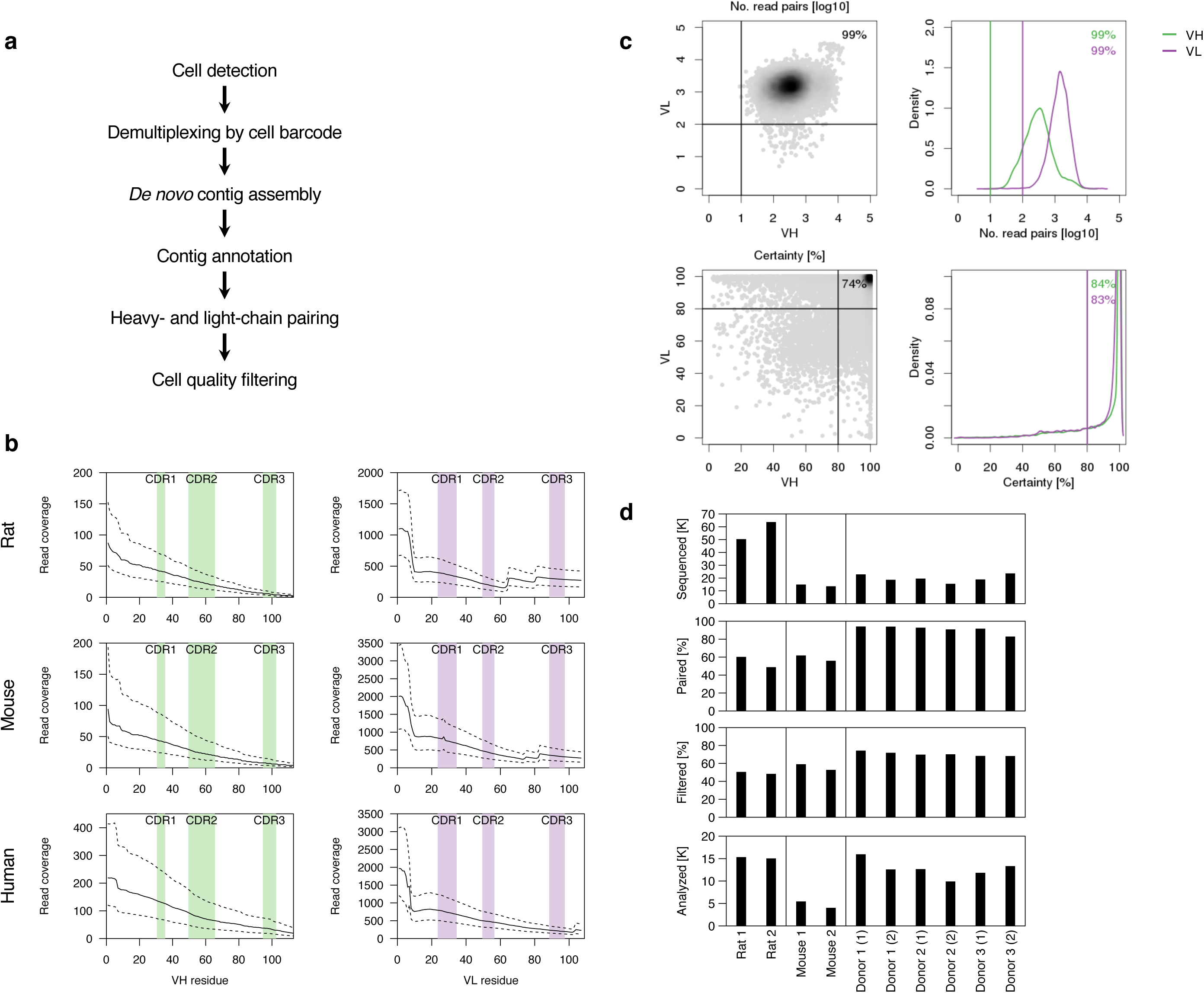
Computational framework for VH-VL pairing. (a) Data analysis workflow. (b) Read coverage for assembled VH (left) and VL (right) contigs for three different species. Positions within VH and VL are based on Kabat numbers, CDR regions are indicated by shaded regions. Solid lines indicate the median, dashed lines interquartile range. (c) Cell quality filtering based on minimum number of reads for VH and VL assemblies (top) and VH and VL certainty (bottom). (d) Overview of B-cell repertoire data generated for this study. Bar graphs show data for independent samples. From top to bottom, number of cells captured, percentage of cells with at least one complete VH and VL assembly, percentage of cells that pass quality filtering, number of cells that pass quality filtering.

**Figure 3.**
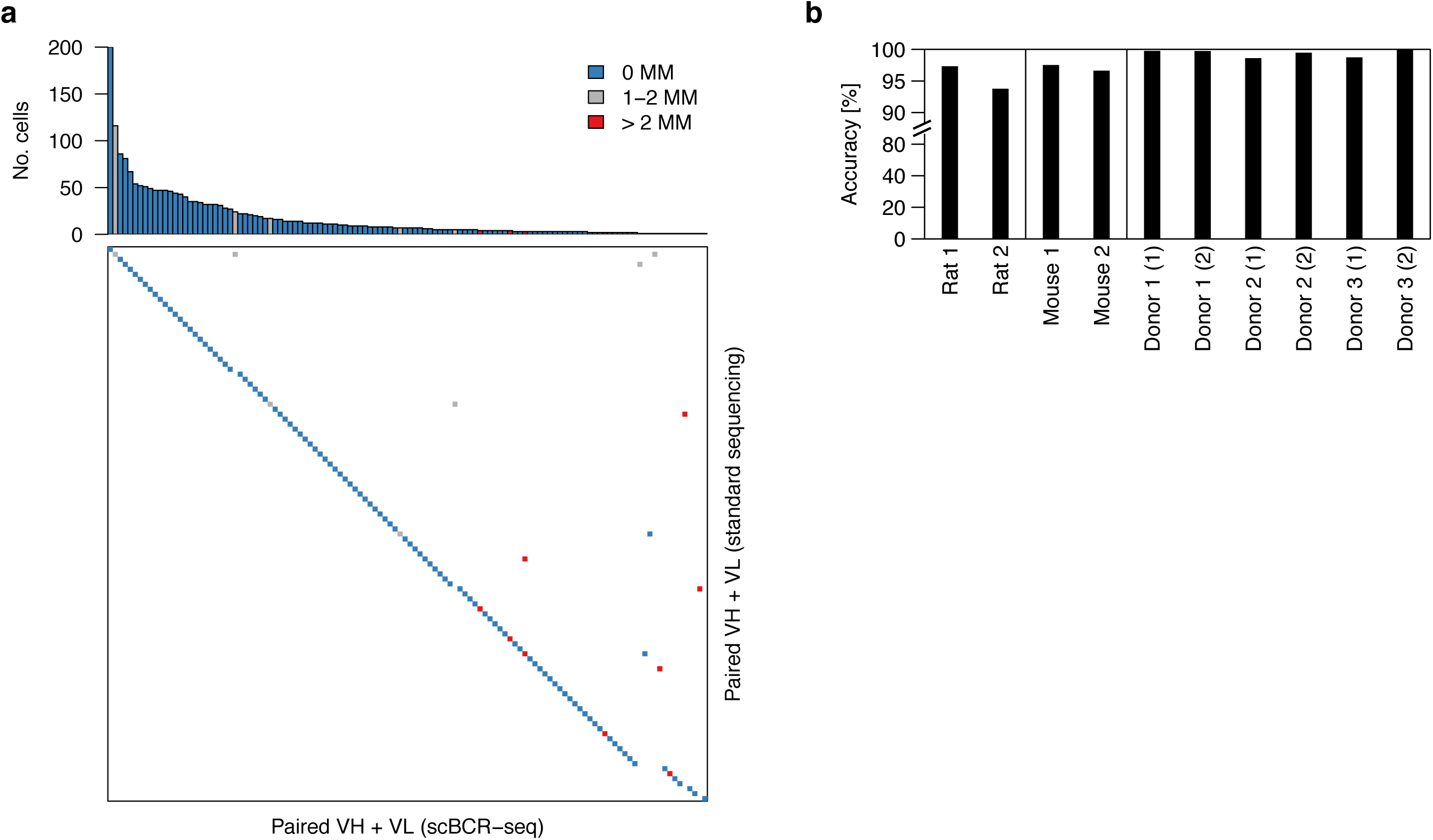
Technology validation. (a) Comparison of unique VH-VL nucleotide sequences obtained by scBCR-seq (columns) and reference VH-VL pairs obtained by a standard sequencing approach (rows). Blue, gray and red boxes indicate VH-VL sequences validated with 0, 1-2 and > 2 mismatches, respectively. Top panel shows number of cells for a particular VH-VL sequence in scBCR-seq data. (b) Pairing accuracy based on V_L_ concordance for VH-VL pairings with identical V_H_ and CDR-H3.

**Figure 4.**
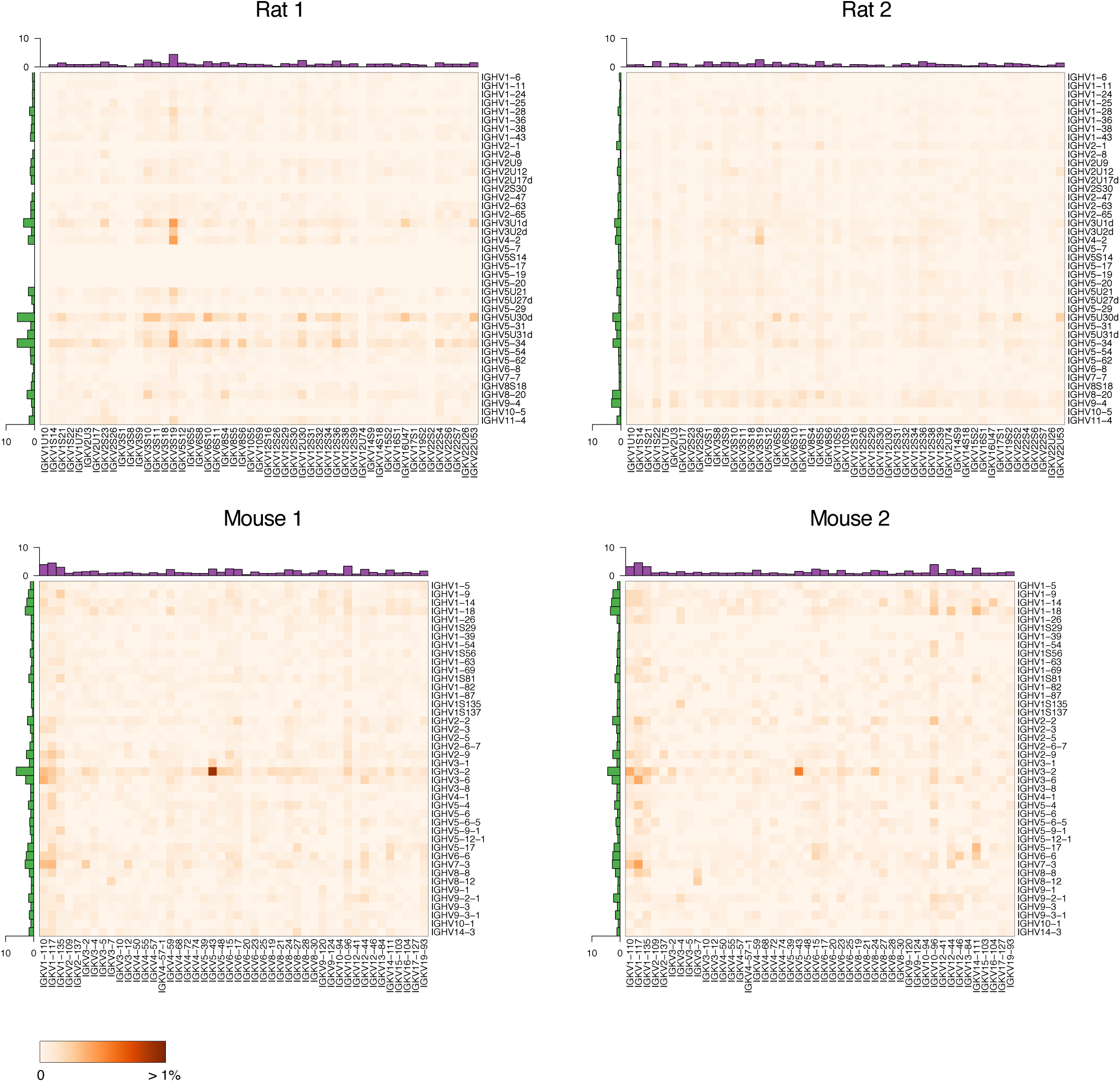
Germline V gene pairing for B-cell repertoires from rats (top) and mice (bottom). Heatmaps show the percentage of lineages with a particular V_H_-V_L_ pairing. Row and column histograms indicate marginal V_H_ and V_L_ frequencies, respectively.

**Figure 5.**
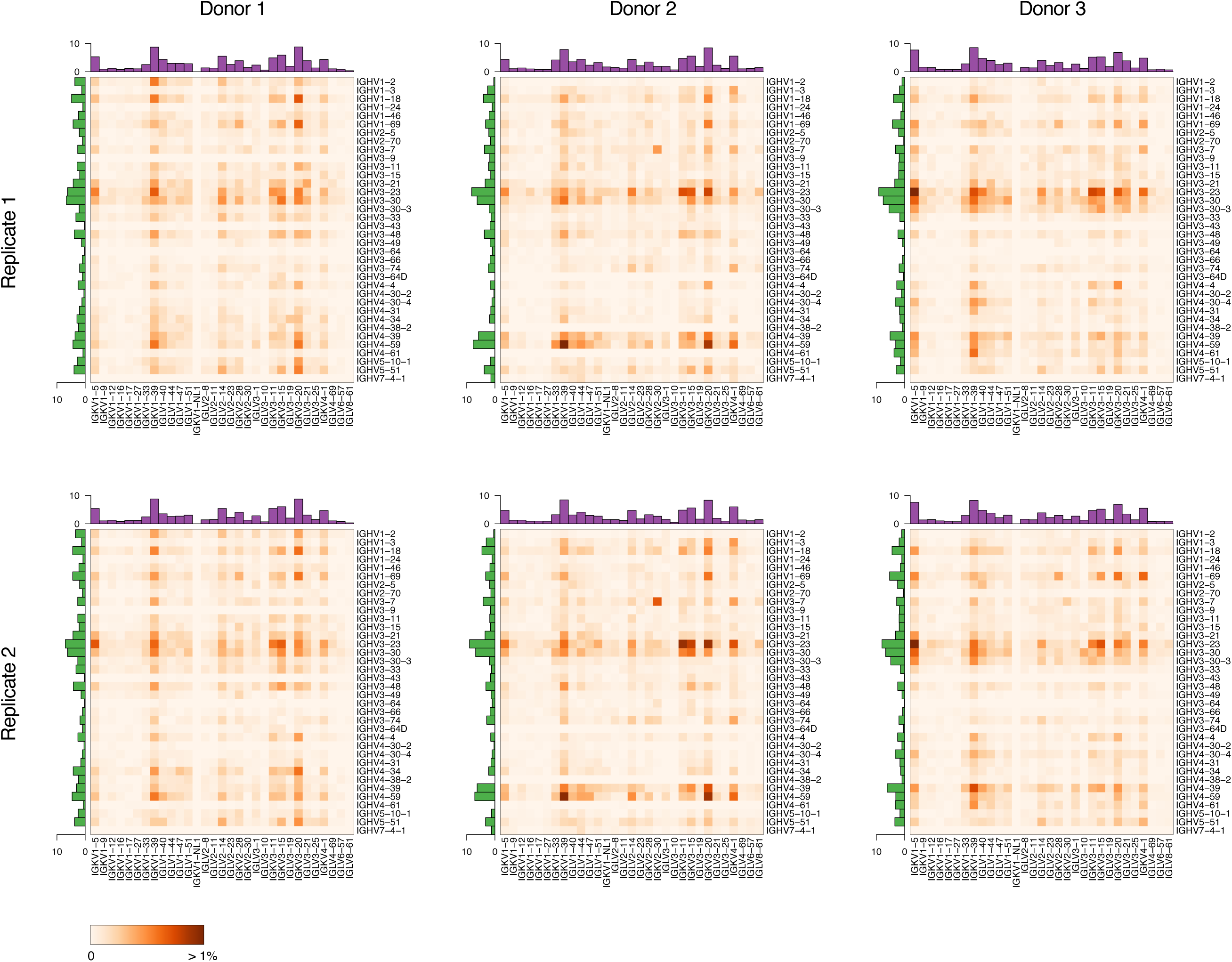
Germline V gene pairing for B-cell repertoires from three human donors. Top and bottom panels are from replicates from the same donor at different time points. Otherwise as in Fig. 4.

### VH-VL pairing accuracy

To assess the accuracy of assembled VH and VL sequences and VH-VL pairing, we used hybridomas obtained after immunizing rats with chicken ovalbumin as a test sample. We analyzed individual clones by a standard sequencing approach to obtain a reference set of paired VH-VL sequences for 127 unique clones (see Methods). Clones were pooled, grown in culture, and subjected to scBCR-seq. scBCR-seq yielded high-quality VH-VL sequence pairs for 1,989 cells, representing 120 unique VH-VL sequence pairs. More than 90% of cells (1,801/1,989) showed a perfect match with one of the reference VH-VL pairs and more than 99% (1,972/1,989) matched one of the reference VH-VL pairs when allowing for up to two nucleotide mismatches (Fig. 3a).

Next, we assessed VH-VL pairing accuracy based on B-cell repertoire data sets from human, mouse and rat samples. Cells within the same B-cell lineage are expected to have light chains with concordant light-chain germline V gene (V_L_) in the majority of cases^8^. We assumed cells with identical heavy-chain V germline (V_H_) gene and CDR-H3 belonged to clonally related B cells, and calculated the percentage of cells with concordant paired V_L_ germline. Concordance in the subset of lineages with more than one B cell was 98% for human (based on 754–2,131 cells in 351–894 lineages), 96% for mouse (445 and 979 cells in 148 and 225 lineages, respectively), and 93% for rat data sets (209 and 598 cells in 97 and 269 lineages, respectively) (Fig. 3b). These estimates are likely lower bounds for pairing accuracy, due to coincidental V_H_ and CDR-H3 matches in some B-cell lineages, in particular for short CDR-H3 sequences more prevalent in rodents^9^

### Rat and mouse B-cell repertoires

We profiled IgM^neg^ B-cell repertoires from two non-immunized rats (n = 30,380) and IgG^pos^ B-cell repertoires from two mice (n = 9,459) (Fig. 2d, Supplementary Fig. 6a,b). Data at single-cell resolution allowed us to characterize unique B-cell lineages and quantify their expansion based on the number of cells observed for each lineage. We defined B-cell lineages by grouping cells with identical V_H_ and V_L_ germline genes and requiring at least 80% nucleotide identity in the CDR-H3 region. For the two rat samples, only 7% (1,081/15,338) and 3% (431/15,042) of cells belonged to clonally expanded lineages (Supplementary Fig. 7). For the two mouse samples, 13% (699/5,448) and 34% (1,353/4,011) of cells belonged to clonally expanded lineages, suggesting some level of antigenic stimulation in these mice (Supplementary Fig. 7). We asked whether the identified B-cell lineages showed preferential usage of particular V_H_ genes, V_L_ genes or V_H_-V_L_ gene pairings. We observed that some germline gene segments were consistently used more frequently than others across replicates (Fig. 4). This was particularly noticeable for mouse V_H_ and V_L_ genes (r = 0.89, r = 0.84, Spearman correlation coefficient) and to a lesser extent for rat V_H_ genes (r = 0.52) (Supplementary Fig. 8). For example, IGHV3-2 was the most commonly used mouse V_H_ gene in both animals, present in 6.2% and 4.4% of lineages, respectively. After correcting for variation in V_H_ and V_L_ gene usage, some individual V_H_-V_L_ gene pairings showed higher than expected frequencies across replicates (Supplementary Fig. 8). For example, both mouse samples showed increased frequencies for IGHV8-12:IGKV3-7 (9 and 7 lineages) and IGHV3-2:IGKV5-43 (45 and 16 lineages). Higher than expected pairings may be due to immune responses to common antigens. Consistent with this interpretation, several of the IGHV8-12:IGKV3-7 lineages showed evidence for expansion in both animals (3/9 and 5/7 lineages, respectively).

**Figure 6.**
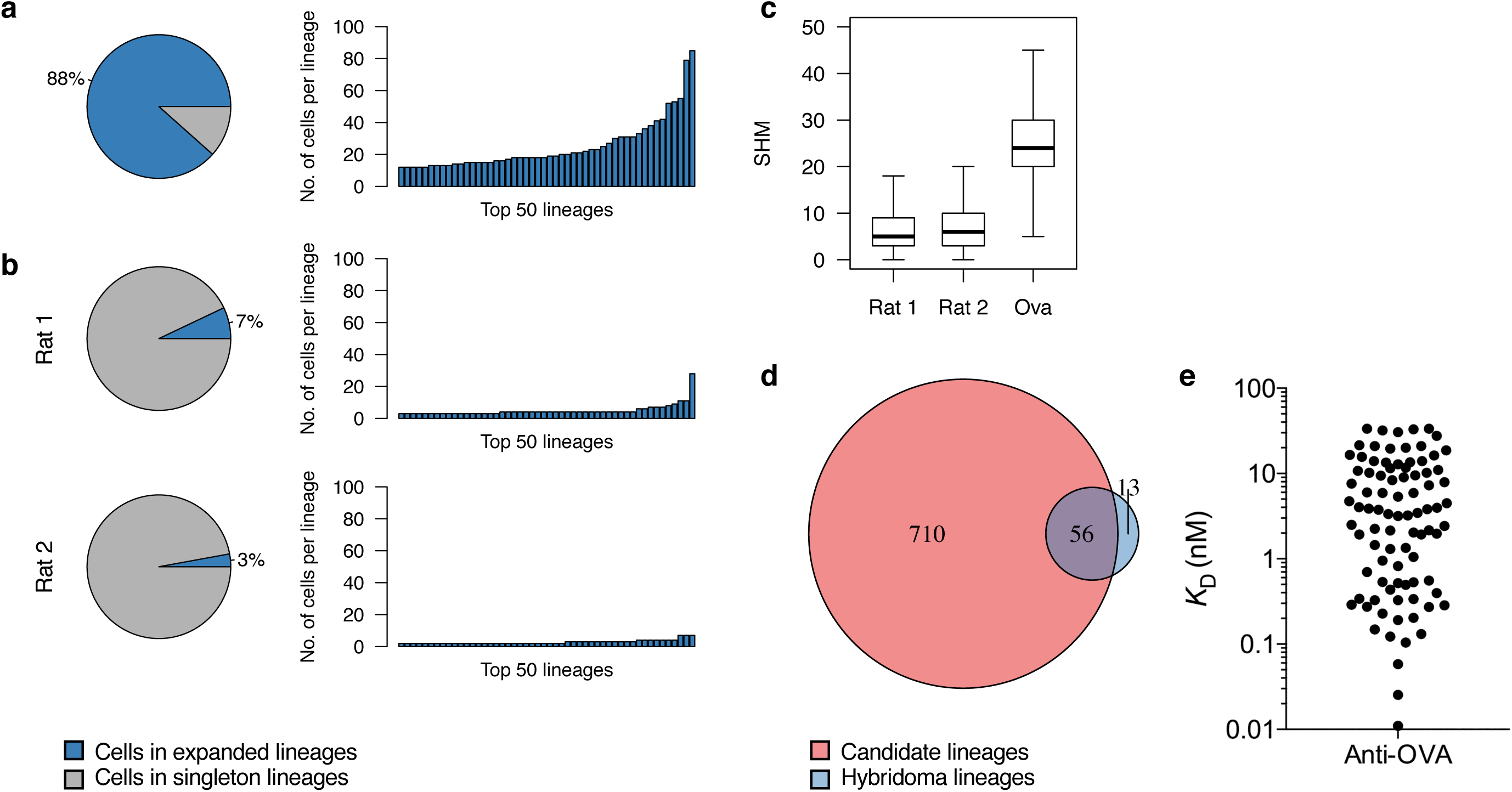
Discovery and validation of antigen-reactive antibodies. (a) Lineage expansions among OVA antigen-reactive B cells. Pie charts indicate percentage of cells belonging to expanded lineages. Bar charts indicate the number of cells for the top 50 lineages. (b) Lineage expansions observed in B-cell repertoires for two non-immunized rats, otherwise as in (b). (c) Somatic hypermutations (SHM) for heavy- and light-chain V germline gene segments for B-cell repertoires from non-immunized rats (Rat 1, Rat 2) and OVA antigen-reactive B cells from immunized rats. Boxes indicate the interquartile range (IQR), center lines the median, whiskers extend to the most extreme data point within 1.5 × IQR from the box. (d) Overlap in lineages identified from direct sequencing of individual antigen-reactive B cells by scBCR-seq (red) and concomitant hybridoma experiment (blue). (e) Validation of candidate OVA antigen-reactive clones. Shown are monovalent affinities of expressed antibodies to OVA.

### Human B-cell repertoires

Next, we analyzed human IgG^pos^ B-cell repertoires from three donors, each profiled at two different time points (n = 76,276) (Fig. 2d, Supplementary Fig. 6c). All samples showed evidence for lineage expansion, with 14–26% of cells belonging to expanded lineages (Supplementary Fig. 7). V_H_ and V_L_ gene usage was highly reproducible between replicates from the same individual (r > 0.98, Supplementary Fig. 9). Overall V_H_ and V_L_ gene usage was similar among donors (r > 0.65 and r > 0.87 for V_H_ and V_L_, respectively, Supplementary Fig. 9) and in general agreement with the known distribution of germline usage in human repertoires^10, 11^. Some donors lacked antibodies for a subset of germline gene segments (e.g. IGHV3-9 in Donor 1 or IGHV4-38-2 in Donor 2), most likely due to genotype differences in the germline repertoires^11^. Interestingly all samples showed higher than expected pairing frequencies for IGHV3-7:IGKV2-30 (19–62 lineages per sample) (Fig. 5, Supplementary Fig. 9). To rule out a technical artifact due to the profiling method, we reanalyzed published V_H_-V_L_ pairing information for naïve and antigen-experienced human B cells that were obtained by overlap extension RT-PCR and independent computational methods^12^. Interestingly, the published data showed strongest enrichment for IGHV3-7:IGKV2-30 among all V germline gene pairings for antigen-experienced, but not naïve B cells, suggesting this pairing may be the result of stereotypical immune responses (Supplementary Fig. 10).

### Single-cell sequencing enables rapid discovery of antigen-reactive antibodies

To assess the potential of scBCR-seq for antibody discovery, we immunized rats with chicken ovalbumin (OVA) and subjected IgM^neg^/OVA^pos^ lymph node B cells from three immunized animals to scBCR-seq (Supplementary Fig. 6d). After quality filtering we obtained VH-VL pairing information for 3,091 B cells (Supplementary Table 12). As expected, significant clonal expansion was inferred in this dataset with 88% of cells belonging to clonally expanded lineages (Fig. 6a). Of 766 unique B-cell lineages, 288 (38%) were represented by three or more individual B cells. By contrast, IgM^neg^ B cells from naïve rats showed limited evidence for clonal expansions (Fig. 6b). Chain pairing accuracy assessed by light-chain germline concordance was 99%, consistent with results obtained for naïve rats. Somatic mutation load in the V_H_ and V_L_-derived regions (i.e. excluding CDR-H3 and J regions in both chains) was higher in anti-OVA cells than in IgM^neg^ B cells from naïve rats (Fig. 6c). In addition to directly sequencing B cells from OVA-immunized animals, we also generated and sequenced OVA-specific hybridomas derived from a fraction of the IgM^neg^ B cells from the same rats. In this data set we identified 69 unique B-cell lineages, 56 of which were shared with those identified by direct B-cell scBCR-seq (Fig. 6d). Thus scBCR-seq recovered 81% (56/69) of anti-OVA lineages from the hybridoma experiment, and identified an additional 710 candidate lineages.

### scBCR-seq identifies multiple antigen-reactive antibodies

We selected a set of 96 BCR (VH-VL) pairs identified from the OVA^pos^ B-cell scBCR-seq data for functional analysis. We selected the BCR sequence pairs from a range of lineage sizes observed in the scBCR-seq data (Supplementary Table 13). A random B-cell clone was selected from lineages represented by 3 or more individual B cells. For lineages with 1–2 cells, which may be enriched for antigen-negative clones due to imperfect cell sorting, we prioritized clones with high read count based on the assumption that these may be antibody-secreting cells induced by an active immune response. Of the 96 selected BCR sequence pairs, 77 were from lineages with three or more cells and 19 were from lineages with 1–2 cells. Of the 96 selected clones, 93 were from lineages not found in the hybridoma panel and 3 had non-identical but clonally related OVA-specific monoclonal antibodies. Synthesized VH and VL DNA sequences were cloned into human IgG1/κ expression vectors and rat/human chimeric IgG was expressed in mammalian cells (see Methods). We found that 93 of the 96 clones showed robust protein expression with a median yield of 85 µg of purified IgG per ml of culture (Supplementary Table 13). All but 4 of the 93 clones (96%) specifically bound OVA in an enzyme-linked immunosorbent assay (ELISA) (Supplementary Fig. 11a). To assess the importance of correct chain pairing for antigen binding in this panel, we shuffled each of the heavy and light chain expression constructs with a non-cognate chain to generate 96 additional antibodies with non-native light and heavy chain pairs. Eighty-six of the 96 shuffled clones expressed as IgG and only 5 of these showed OVA reactivity (Supplementary Fig. 11b), confirming the relevance of accurate chain pairing for identification of antigen-binding antibodies. We further determined the affinity of the 93 expressed antibodies for OVA by surface plasmon resonance (SPR).

Eighty-nine clones, including 3 clones with weak binding in ELISA, bound to OVA in SPR, with monovalent equilibrium dissociation constants ranging from the limit of detection (10 pM) to 30 nM (Fig. 6e, Supplementary Fig. 11, Supplementary Table 13). Interestingly, binding affinity did not correlate with lineage size or read counts per cell (Supplementary Table 13). Overall, 86 of the 93 expressed antibodies (92%) bound OVA in both ELISA and SPR, and 92 antibodies (99%) bound OVA in one of the two assays. These findings show scBCR-seq to be a robust tool for rapid discovery of a large panel of antigen-reactive antibodies.

## DISCUSSION

Antibody discovery methods based on molecular cloning of antibody repertoires are limited by the inability to easily process large numbers of single cells in parallel. Next-generation sequencing technologies that capture paired antibody heavy- and light-chain information have been described but require custom equipment and do not yield full-length variable region sequences or true single-cell sequence information. We show that scBCR-seq yields full-length antibody variable region sequences from a variety of species at an individual cell level. While framework region 4 sequences were partially truncated, the short missing regions, unlike CDR regions directly involved in antigen binding, can be easily reconstructed from germline sequences with minimal impact on antigen binding properties. Importantly, scBCR-seq does not require a complex set of primers covering the highly diverse V_H_ and V_L_ germlines of human and rodent repertoires and somatic mutations, minimizing biases and blind spots introduced in the amplification process. Thus scBCR-seq should be applicable with minimal modification to other species, including those for which only constant region sequences are known.

Our results show that scBCR-seq can be used for rapid discovery of large, diverse panels of high-affinity antigen-specific antibodies with natively paired heavy- and light-chains when combined with high-quality antigen-specific B cells sorting. This is supported by the high rate of antigen binders and the high overlap between the anti-OVA B-cell repertoire and hybridoma sequencing. The highly consistent V_L_ germline pairing with clonally related heavy chains support the high quality of chain pairing information yielded. Here we only tested a subset of the diversity yielded by scBCR-seq. We expect that more in-depth analyses of clones within expanded antigen-specific B-cell lineages will allow easy and rapid identification of higher affinity variants within each lineage. In addition, the availability of datasets with thousands of paired-chain, full-length variable region sequences from individual antigen-specific B cells will allow detailed large-scale characterization of the dynamics of immune responses, and the functional impact of germline usage, clonal expansion and somatic mutation on antibody repertoires.

## ACKNOWLEDGMENTS

We thank the Genentech NGS Lab for their support during the course of this project. We also thank Qing Zhang for her support during the development of the Absolve annotation pipeline.

## AUTHOR CONTRIBUTIONS

Conceptualization LDG, IH, SS; Formal analysis: LDG, IH; Investigation: YJC, JW, SC, YH, KS, ZL, BSJ, AA, KBP, ZM; Methodology: LDG, YJC, KHH, SG, BSJ, JS, ZM, IH, SS; Software: LDG, SG; Supervision: JS, DS, ZM, IH, SS; Writing – original draft: LDG, IH, SS; Writing – review & editing: All authors. Based on CRediT contributor taxonomy.

## COMPETING INTERESTS

The authors are employees of Genentech or SciGenom Labs. Genentech authors hold shares in Roche.

## METHODS

### Isolation of IgG^pos^ B cells from human blood

Blood was obtained from healthy human donors before and on days 6 and 7 after vaccine immunization with the seasonal influenza Fluzone (year 2018). Samples were obtained after written informed consent was provided and ethical approval granted from the Western Institutional Review Board. Blood samples were diluted in phosphate-buffered saline (PBS) at a 1:1 (volume/volume) ratio and layered on top of a Ficoll-Paque-PLUS medium cushion (density 1.077 g/ml, GE Healthcare Life Sciences). Samples were centrifuged for 30 minutes at room temperature at 400 g with a soft stop. The interface layer containing peripheral blood mononuclear cells was collected and resuspended in FACS staining buffer (PBS, 0.5% BSA, 2 mM EDTA) for marker staining and cell sorting. Single-cell suspensions at 5 × 10^7^ cells per ml were stained with a cocktail of fluorochrome conjugated human antibodies (BD Biosciences, San Jose, CA), anti-human IgG APC, anti-human CD20 PE Cy7, anti-human CD4 APC Cy7, Propidium iodide (PI) P4864 (Sigma-Aldrich, St. Louis, MO). IgG^pos^ B cells were enriched using PE-dump channel cocktail antibodies to exclude granulocyte, monocyte/macrophage, dendritic, NK, CD8 cell population. IgG^pos^ B cells (100,000 cells from each donor) were sorted on a FACSAria II cell sorter (BD Biosciences, San Jose, CA) and collected for scBCR-seq (Supplementary Fig. 12).

### Preparation of naïve Balb/c mouse and rat lymph node (LN) B cells

All animals used in this study were housed and maintained at Genentech in accordance with American Association of Laboratory Animal Care guidelines. All experimental studies were conducted under protocols approved by the Institutional Animal Care and Use Committee of Genentech Lab Animal Research in an Association for Assessment and Accreditation of Laboratory Animal Care International-accredited facility in accordance with the Guide for the Care and Use of Laboratory Animals and applicable laws and regulations. Axillary brachial, mesenteric, inguinal, iliac and popliteal LNs were collected from naïve Balb/c mice or Sprague Dawley (SD) rats. Single-cell suspensions were prepared by crushing LNs through 70 µm polyester mesh sterile cell strainers (6 Netwell inserts plate, Corning). Murine B cells were enriched by negative selection of IgM^pos^ cells with a mouse B cell isolation kit (Miltenyi Biotec, #130-090-862, Bergisch Gladbach, Germany), resuspended in FACS staining buffer at 5 × 10^7^ cells per ml, stained with a cocktail of fluorochrome conjugated anti-mouse B220 APC (BD Biosciences, San Jose, CA) anti-mouse IgG FITC (Bethyl Laboratories, Montgomery, TX) for sorting of IgG^pos^ B cells (100,000 cells) on a FACSAria II cell sorter (Supplementary Fig. 13). Rat B cells were enriched using a cocktail of biotinylated anti-rat CD4 (Clone OX-35), biotinylated anti-rat CD8a (Clone OX-8), biotinylated anti-rat 11b/c (Clone OX42), biotinylated anti-rat CD161 (Clone 10/78) and biotinylated anti-rat granulocyte marker (Clone HIS48) antibodies (BD Biosciences, San Jose, CA) followed by magnetic separation (Miltenyi Biotec, San Diego, CA) using streptavidin beads, stained with anti-rat IgM (Clone G53-238, BD Biosciences, San Jose, CA) conjugated to PE Cy7, CD45RA-APC Cy7, a dump cocktail of antibodies (BV510 anti rat granulocyte (Clone HIS48, BD Biosciences, San Jose, CA), BV510 anti rat 11b/c (Clone OX42, BD Biosciences, San Jose, CA), BV510 anti rat CD161a (Clone 10/78, BD Bioscience, San Jose, CA), and BV510 anti rat CD4 (Clone OX35, BD Bioscience, San Jose, CA)) and sorted for B cells (100,000 cells) on a FACSAria II cell sorter (Supplementary Fig. 14).

### Immunization of rats and antigen-specific cell sorting

Three SD rats (Charles River, Hollister, CA) were immunized with 100 µg OVA (Sigma-Aldrich, St. Louis, MO) in Complete Freund’s adjuvant (BD, Franklin Lakes, NJ) followed by bi-weekly boosts of 50 µg OVA in incomplete Freund’s adjuvant divided in three sites (intraperitoneally, subcutaneously at base of tail, at nape of neck and in both hocks). Multiple lymph nodes were harvested from each rat two days after the last immunization, pooled and enriched for B cells as described above. Half of the enriched B cells were stained with anti-rat IgM (Clone G53-238, BD Biosciences, San Jose, CA) conjugated to PE Cy7, APC-labeled OVA conjugated to Alexa Fluor 647 (Thermo Fisher Scientific, Waltham, MA), CD8-per CPCy5.5, CD45RA-APC Cy7, a dump cocktail of antibodies (BV510 anti rat granulocyte (Clone HIS48, BD Biosciences, San Jose, CA), BV510 anti rat 11b/c (Clone OX42, BD Biosciences, San Jose, CA), BV510 anti rat CD161a (Clone 10/78, BD Bioscience, San Jose, CA), and BV510 anti rat CD4 (Clone OX35, BD Bioscience, San Jose, CA)) and sorted for dump channel^neg^/CD8^neg^/CD45RA^pos^/IgM^neg^/OVA^pos^ cells in a FACSAriaIII sorter (BD, Franklin Lakes, NJ) (Supplementary Fig. 15). A total of 32,845 such IgM^neg^/OVA^pos^ events were detected from the three rats and processed for scBCR-seq. The other half of the B cells were used for hybridoma generation. IgM-negative B cells were purified from lymphocytes using biotinylated anti-rat IgM (Clone G53-238, BD Biosciences, San Jose, CA) and magnetic separation (Miltenyi Biotec, San Diego, CA) using streptavidin beads followed by fusion with Sp2ab mouse myeloma cells (Abeome, Athens, GA) via electrofusion (Harvard Apparatus, Holliston, MA). Fused cells were incubated at 37°C, 7% CO_2_, overnight in Clonacell-HY Medium C (StemCell Technologies, Vancouver, BC, Canada), before centrifugation and resuspension in Clonacell-HY Medium E (StemCell Technologies, Vancouver, BC, Canada) supplemented with HAT (Sigma-Aldrich, St. Louis, MO) and plating into 12-well plates and allowed to grow at 37°C, 7% CO_2_. Four days after plating, hybridomas were stained with anti-rat IgG (goat polyclonal, Jackson ImmunoResearch, West Grove, PA) conjugated to Alexa 488 and APC-labeled OVA conjugated to Alexa Fluor 647 (Thermo Fisher Scientific, Waltham, MA) and sorted for IgG^pos^/OVA^pos^ cells in a FACSAriaIII sorter (BD, Franklin Lakes, NJ) (Supplementary Fig. 16). A total of 649 IgG^pos^/OVA^pos^ hybridoma cells were individually deposited into 96-well plates containing Medium E (StemCell Technologies, Vancouver, BC, Canada). Supernatants were screened by ELISA against OVA seven days after sorting. A total of 379 clones producing antibodies binding to OVA in ELISA were pooled and submitted to scBCR-seq. ELISA-positive hybridoma clones were also individually sequenced as previously described, leading to the identification of 127 unique clones^13^.

### 10× single-cell B-cell receptor library construction and sequencing

Sample processing for single B cell receptor (BCR) V(D)J clonotype was done using Chromium Single Cell 5’ Library and Gel Bead Kit following the manufacturer’s user guide (10x Genomics, Pleasanton, CA, CG000086_SingleCellVDJReagentKitsUserGuide_RevB). After FACS sorting, cells were spun down, resuspended in 3% fetal bovine serum (Sigma-Aldrich, St. Louis, MO)/phosphate buffer solution (Thermo Fisher Scientific, Waltham, MA) and subjected to cell quality control using Vi-CELL XR cell counter (Beckman Coulter, Brea, CA). All of the processed B cells had cell viability >90%. After determining cell density, cells were injected into 8 channels for rat samples, and 4 channels for mouse and human samples, aiming to achieve ∼6,000 cells per channel. Gel Beads-in-Emulsion (GEMs) were formed in channels of a chip in the 10 × Chromium instrument, and then collected into an Eppendorf plate for GEM reverse transcription (GEM-RT) reaction. After GEM clean-up, GEM-RT products were subjected to two rounds of 14 PCR cycles using custom primers for rat and mouse, and 10× human BCR primers (10 × Genomics, Pleasanton, CA) for human, followed by SPRIselect (Beckman Coulter, Brea, CA) beads clean-up. The 10 × human BCR primers cover human heavy chain isotypes and both κ and λ light chains. Mouse and rat λ chains, which comprise a relatively minor fraction of the repertoire in rodents, were not covered by custom primers (Supplementary Information). Single-cell BCR V(D)J Libraries were prepared following the manufacturer’s user guide (10x Genomics, Pleasanton, CA, CG000086_SingleCellVDJReagentKitsUserGuide_RevB), and profiled using Bioanalyzer High Sensitivity DNA kit (Agilent Technologies, Santa Clara, CA) and quantified with Kapa Library Quantification Kit (Kapa Biosystems, Wilmington, MA). Libraries were sequenced by paired-end sequencing (2 × 150bp) on an Illumina HiSeq2500 or HiSeq4000 (Illumina, San Diego, CA). BCL data were converted to demultiplexed FASTQ files using Illumina bcl2fastq 2.20.

### Cell count estimate

For each library we tallied the number of reads for expected cell barcode sequences. Barcodes from cell-containing droplets were identified as those with read count exceeding a library-specific cutoff, defined as 0.1 times the read count for barcodes at rank 150, the 97.5th percentile for 6,000 targeted cells.

### Contig assembly

We trimmed the first 39 bases for read 1 covering the 16 nt cell barcode, 10 nt unique molecular identifier (UMI) and 13 nt switch oligo. Barcode and UMI sequences were retained for each read. Reads were demultiplexed based on perfect matches to one of the expected barcode sequences. Subsequent processing was done independently for each barcode. If reads for a given barcode exceeded 100,000, they were downsampled to 100,000. Reads were used as input for de novo assembly with SSAKE (3.8.5)^14^ with options ‘-p 1 -c 1 -w 1 -e 1.5’ and expected mean insert size 600. We defined concordant read pairs as those with both reads fully embedded in the assembly and paired in the expected orientation. Contigs without concordant read pairs were discarded. Contigs were trimmed to regions supported by concordant read pairs. In cases with more than one contiguous supported region, the region with highest number of concordant read pairs was selected.

### Contig annotation

Sequences were annotated with a custom bioinformatics pipeline for variable domain analysis (https://github.com/Genentech/Absolve). Sequences were aligned using HMMER (http://www.hmmer.org/) to Hidden Markov Models that were trained on heavy and light germline amino acid sequences from the International Immunogenetics (IMGT) database^15^ in order to determine chain identity, framework and CDR boundaries and residue Kabat numbering^16^. Sequence germline assignments were determined by aligning to IMGT heavy and light V and J germline database sequences^17^ using the SSW library^18^ and selecting the highest scoring germline. In order to assess the fidelity of germline classifications, Absolve was benchmarked with a set of 2,000 simulated VH sequences, including random combinations of 252 V, 44 D and 13 J human segments and alleles and random trimming and addition of nucleotides. Each sequence was mutated up to 40 times, preferentially in CDR regions, to yield a set of 80,000 in-frame VH sequences with a variable number of mutations without stop codons. Processing of the simulated dataset with Absolve and IgBlast^19^ yielded comparable V_H_ and J_H_ germline call error rates (< 0.5% in V_H_ at the highest mutational load). Reference germlines were from the IGMT database except for 52 additional rat germlines mined from *Rattus norvegicus* whole genome contigs from GenBank and germline sequences deduced from SD rat repertoire sequencing data (I. Hötzel, unpublished) (Supplementary Information).

### Heavy- and light-chain pairing

We considered contigs identified as either heavy-or light-chain variable domains with HMM score ≥ 30. Identified variable domains were considered complete if they contained all of framework region 1 (FW1) and the first four positions of framework region 4 (FW4). Amino acid sequences downstream of FW4 position 4 were ignored. For each cell we reported the complete heavy- and light-chain variable domain with highest number of concordant read pairs. For each reported heavy- and light-chain variable domain we calculated a certainty score, defined as the number of read pairs supporting the top contig divided by the total number of read pairs for all identified heavy- and light-chain contigs, respectively. Filtered high-quality pairings are those with (1) top heavy chain supported by ≥ 10 read pairs (2) top light chain supported by ≥ 100 read pairs, and (3) certainty ≥ 80% for both heavy and light chain.

### Lineage definition

Filtered cells were grouped into lineage clusters. Two cells were grouped together if they had identical germline V_H_ and V_L_ genes, ignoring allele number, identical CDR-H3 length and ≥ 80% identical CDR-H3 sequence. J_H_ and J_L_ genes were ignored for lineage definition due to the lower sequence coverage in this region.

### Antibody expression and binding assays

Selected B-cell clones from the OVA immunization dataset were produced by DNA synthesis and cloned in mammalian expression vectors^20^ as chimeric human IgG1/kappa antibodies. Sequences with clearly incorrect FR4 terminal sequences were corrected based on the assigned J germline information for that clone but no additional coding changes were introduced in sequences. Antibodies were transfected into Expi293 cells at 1 ml scale and purified in batch mode by protein A chromatography^21, 22^. Antibodies with mispaired chains were made by combining each of the 96 heavy chain clones with a light chain clone from a different anti-OVA antibody. Purified IgGs were tested for binding to OVA in ELISA and in an SPR assay using a BIAcore T200 apparatus (GE Life Sciences, Piscataway, NJ) in a protein A capture format. Soluble monomeric OVA was the analyte in SPR, using the single cycle kinetics method at 25°C.

## Data availability

Raw sequence data generated during the current study will be made available in the Gene Expression Omnibus (GEO) repository before publication.

## Code availability

The custom bioinformatics pipeline for variable domain analysis is available under the MIT License on GitHub (https://github.com/Genentech/Absolve).

## Supplementary Figures

**Supplementary Figure 1**

Schematic of single-cell cDNA generation and library construction following the manufacturer’s user guide (10x Genomics, Pleasanton, CA, CG000086_SingleCellVDJReagentKitsUserGuide_RevB). Only heavy chain is shown.

**Supplementary Figure 2**

Custom PCR primers targeting the rat and mouse heavy- and light-chain constant regions. (a) Multiple sequence alignment of rat heavy-chain isotype constant regions. (b) Rat κ light-chain constant region. (c) Multiple sequence alignment of mouse heavy-chain isotype constant regions. (d) Mouse κ light-chain constant region.

**Supplementary Figure 3**

Cell detection. Number of reads per barcode plotted against barcode rank for each library, barcodes for detected cells are colored in green.

**Supplementary Figure 4**

Read coverage for assembled VH and VL contigs. Otherwise as in Fig. 2b.

**Supplementary Figure 5**

Cell quality filtering. Otherwise as in Fig. 2c.

**Supplementary Figure 6**

Representative B cell sorting gates for naïve mice (a), naïve rat (b), human (c) and pooled OVA-immunized rats (d).

**Supplementary Figure 7**

Lineage expansions. Otherwise as in Fig. 6a,b.

**Supplementary Figure 8**

Pairwise scatter plots of V_H_ frequencies (left), V_L_ frequencies (center), and standardized residuals for V_H_-V_L_ pairings frequencies (right) for two rats (top) and two mice (bottom). Frequencies indicate the percentage of lineages with a particular germline gene segment. IGHV8-12:IGKV3-7 and IGHV3-2:IGKV5-43 are highlighted in red.

**Supplementary Figure 9**

Pairwise scatter plots of V_H_ frequencies (left), V_L_ frequencies (center), and standardized residuals for V_H_-V_L_ pairing frequencies (right) for three human donors profiled in duplicate. Frequencies indicate the percentage of lineages with a particular germline gene segment. IGHV3-7:IGKV2-30 is highlighted in red.

**Supplementary Figure 10**

QQ-plots of standardized residuals for V_H_-V_L_ pairing frequencies for three human donors profiled in duplicate (top) and published data for naïve and antigen-experienced human B cells^12^ (bottom).

**Supplementary Figure 11**

ELISA results for predicted OVA antigen-reactive B cell clones (a) and negative controls obtained by mispairing VH and VL chains (b). Clones are ranked by OVA ELISA signal separately in (a) and (b).

**Supplementary Figure 12**

Human IgG^pos^ B cell sort gating strategy. 1) FSC vs SSC to gate lymphocytes, 2/3) SSC-H/SSC-W, FSC-H/FSC-W gates to exclude cell doublets, 4) FSC/PI for dead cell exclusion (PI-negative gate), 5) IgG-APC/Dump-PE (CD11b, CD11c, CD14, CD16, CD56, CD64, CD8) to exclude PE-positive non-B cells, 6) CD20-PE Cy7/CD4 APC-Cy7 to exclude CD4+ cells, 7) CD20-PECy7/IgG-APC to sort for IgG^pos^ B cells.

**Supplementary Figure 13**

Naïve Balb/c mouse B cell sort gating strategy. 1) FSC vs SSC to gate lymphocytes, 2/3) SSC-H/SSC-W, FSC-H/FSC-W gates to exclude cell doublets, 4) FSC/PI for dead cell exclusion (PI-negative gate), 5) IgG FITC/B220-APC to sort for IgG^pos^ B cells.

**Supplementary Figure 14**

Naïve SD rat B cell sort gating strategy. 1) FSC vs SSC to gate lymphocytes, 2/3) SSC-H/SSC-W, FSC-H/FSC-W gates to exclude cell doublets, 4) FSC/PI for dead cell exclusion (PI-negative gate), 5)CD8-PerCP Cy5.5/Dump-BV510 (CD4, CD11b, CD161a, granulocyte marker) to exclude CD8+BV510+ non-B cells, 6) IgM-PECy7/CD45RA-APC Cy7 to sort for B cells.

**Supplementary Figure 15**

Immunized SD rat, OVA^pos^IgM^neg^ B cell sort gating strategy. 1) FSC vs SSC to gate lymphocytes, 2/3) SSC-H/SSC-W, FSC-H/FSC-W to to exclude cell doublets, 4) FSC/PI for dead cell exclusion (PI-negative gate), 5) CD8-PerCP Cy5.5/Dump-BV510 (CD4, CD11b, CD161a, granulocyte marker) to exclude BV510+ non-B cells, 6) CD45R-APC Cy7/CD8-PerCP Cy5.5 to exclude CD8+ cells, 7) IgM-PE Cy7/CD45R-APC Cy7 to gate for IgM^neg^ B cells, 8) CD45R-APC Cy7/OVA-APC to sort for OVA^pos^IgM^neg^ B cells.

**Supplementary Figure 16**

Immunized SD rat, OVA^pos^IgG^pos^ hybridoma sort gating strategy. 1) FSC vs SSC to gate hybridoma cells, 2/3) SSC-H/SSC-W, FSC-H/FSC-W gates to exclude cell doublets, 4) FSC/PI for dead cell exclusion (PI-negative gate), 5) IgG FITC/BV421 (empty channel) to gate for IgG^pos^ hybridomas, 6)PE (empty channel)/OVA-APC to sort for OVA^pos^IgG^pos^ hybridomas.

## REFERENCES

1. Schroeder, H.W., Jr. Similarity and divergence in the development and expression of the mouse and human antibody repertoires. Dev Comp Immunol 30, 119–135 (2006).

2. Schroeder, H.W., Jr. & Cavacini, L. Structure and function of immunoglobulins. J Allergy Clin Immunol 125, S41–52 (2010).

3. Watson, C.T. & Breden, F. The immunoglobulin heavy chain locus: genetic variation, missing data, and implications for human disease. Genes Immun 13, 363–373 (2012).

4. Reddy, S.T. et al. Monoclonal antibodies isolated without screening by analyzing the variable-gene repertoire of plasma cells. Nat Biotechnol 28, 965–969 (2010).

5. DeKosky, B.J. et al. High-throughput sequencing of the paired human immunoglobulin heavy and light chain repertoire. Nat Biotechnol 31, 166–169 (2013).

6. DeKosky, B.J. et al. In-depth determination and analysis of the human paired heavy- and light-chain antibody repertoire. Nat Med 21, 86–91 (2015).

7. McDaniel, J.R., DeKosky, B.J., Tanno, H., Ellington, A.D. & Georgiou, G. Ultra-highthroughput sequencing of the immune receptor repertoire from millions of lymphocytes. Nat Protoc 11, 429–442 (2016).

8. Hershberg, U. & Luning Prak, E.T. The analysis of clonal expansions in normal and autoimmune B cell repertoires. Philos Trans R Soc Lond B Biol Sci 370 (2015).

9. Zemlin, M. et al. Expressed murine and human CDR-H3 intervals of equal length exhibit distinct repertoires that differ in their amino acid composition and predicted range of structures. J Mol Biol 334, 733–749 (2003).

10. Glanville, J. et al. Precise determination of the diversity of a combinatorial antibody library gives insight into the human immunoglobulin repertoire. Proc Natl Acad Sci U S A 106, 20216–20221 (2009).

11. Boyd, S.D. et al. Individual variation in the germline Ig gene repertoire inferred from variable region gene rearrangements. J Immunol 184, 6986–6992 (2010).

12. DeKosky, B.J. et al. Large-scale sequence and structural comparisons of human naive and antigen-experienced antibody repertoires. Proc Natl Acad Sci U S A 113, E2636–2645 (2016).

13. Chen, Y. et al. Barcoded sequencing workflow for high throughput digitization of hybridoma antibody variable domain sequences. J Immunol Methods 455, 88–94 (2018).

14. Warren, R.L., Sutton, G.G., Jones, S.J. & Holt, R.A. Assembling millions of short DNA sequences using SSAKE. Bioinformatics 23, 500–501 (2007).

15. Lefranc, M.P. IMGT, the international ImMunoGeneTics database: a high-quality information system for comparative immunogenetics and immunology. Dev Comp Immunol 26, 697–705 (2002).

16. Kabat, E.A., Wu, T.T., Perry, H.M., Gottesman, K.S. & Foeller, C. in NIH Publication (1991).

17. Lefranc, M.P., Ehrenmann, F., Ginestoux, C., Giudicelli, V. & Duroux, P. Use of IMGT((R)) databases and tools for antibody engineering and humanization. Methods Mol Biol 907, 3–37 (2012).

18. Zhao, M., Lee, W.P., Garrison, E.P. & Marth, G.T. SSW library: an SIMD Smith-Waterman C/C++ library for use in genomic applications. PLoS One 8, e82138 (2013).

19. Ye, J., Ma, N., Madden, T.L. & Ostell, J.M. IgBLAST: an immunoglobulin variable domain sequence analysis tool. Nucleic Acids Res 41, W34–40 (2013).

20. Eaton, D.L. et al. Construction and characterization of an active factor VIII variant lacking the central one-third of the molecule. Biochemistry 25, 8343–8347 (1986).

21. Luan, P. et al. Automated high throughput microscale antibody purification workflows for accelerating antibody discovery. MAbs 10, 624–635 (2018).

22. Bos, A.B. et al. Optimization and automation of an end-to-end high throughput microscale transient protein production process. Biotechnol Bioeng 112, 1832–1842 (2015).

