## Supplementary Figures for "Massively parallel single-cell B-cell receptor sequencing enables rapid discovery of diverse antigen-reactive antibodies"

#### Supplementary Figure 1

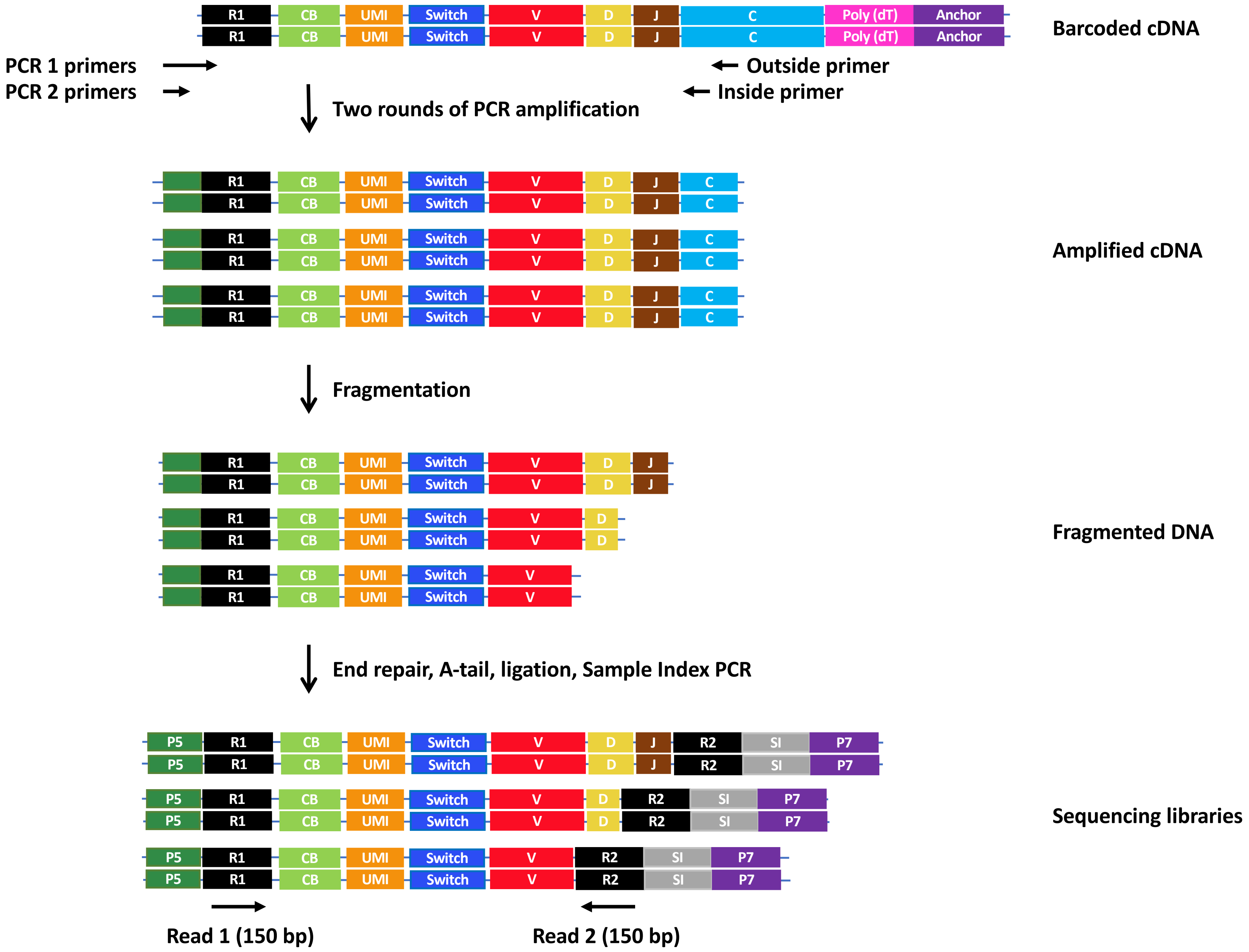

### Supplementary Figure 2

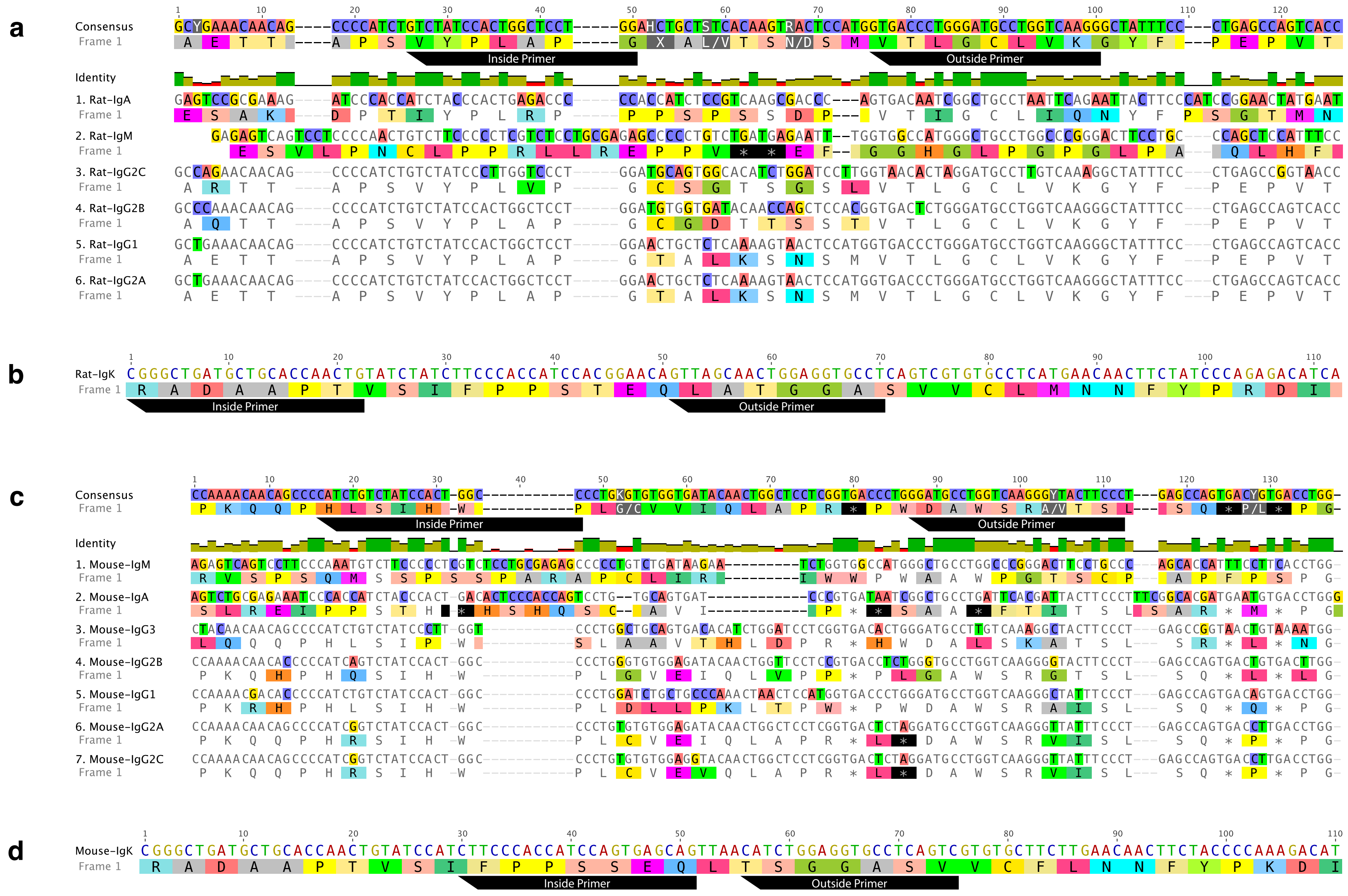

### Supplementary Figure 3

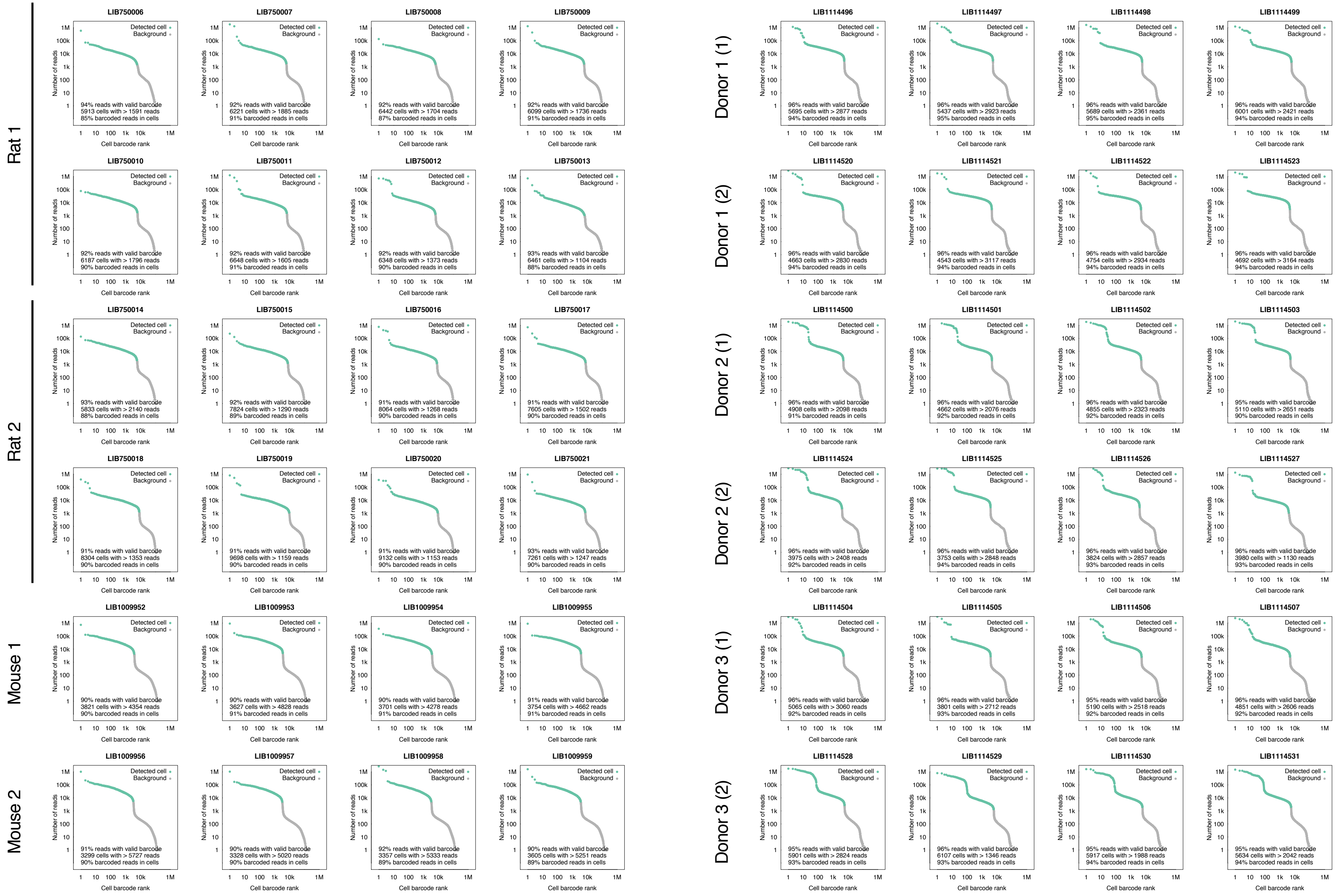

### Supplementary Figure 4

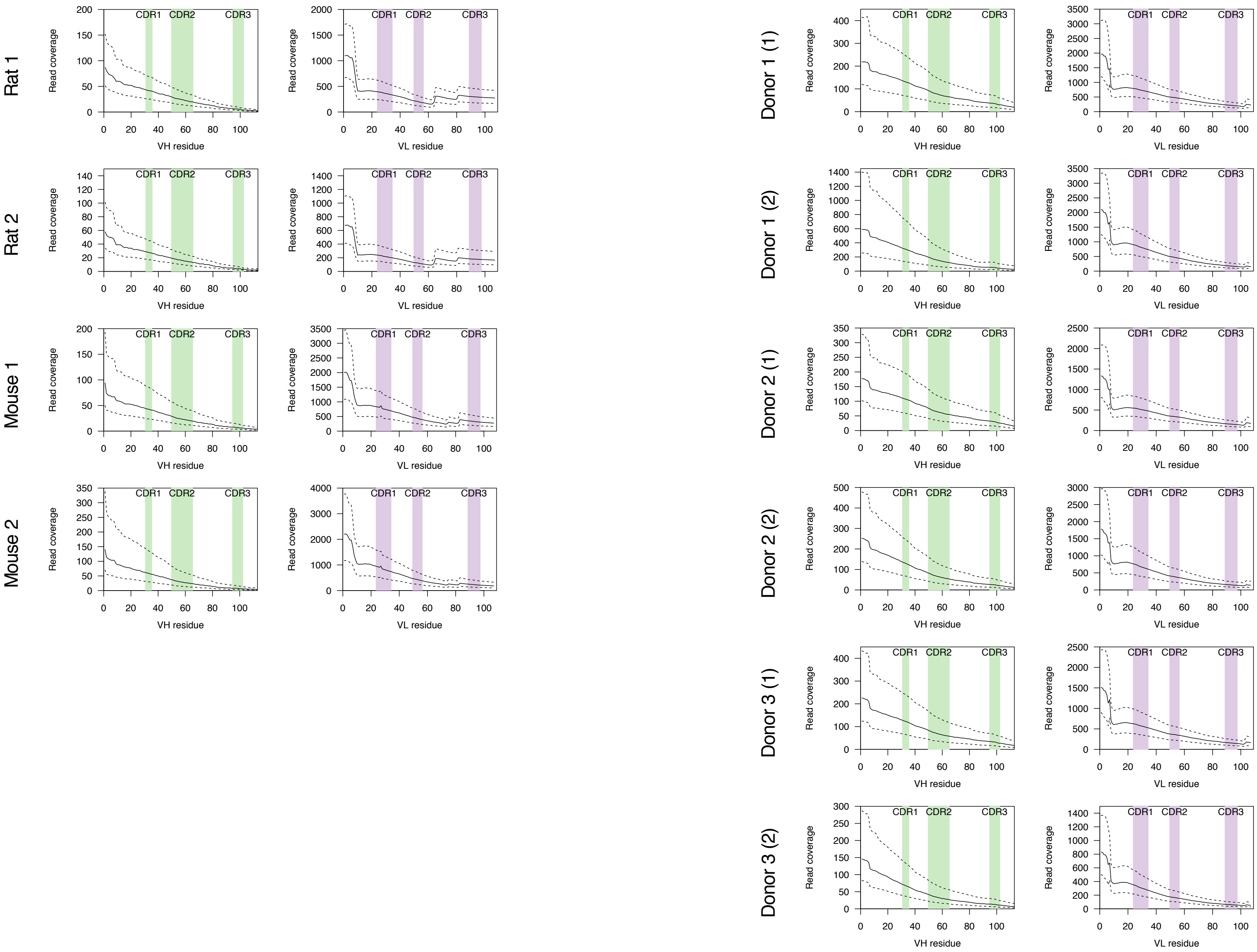

### Supplementary Figure 5

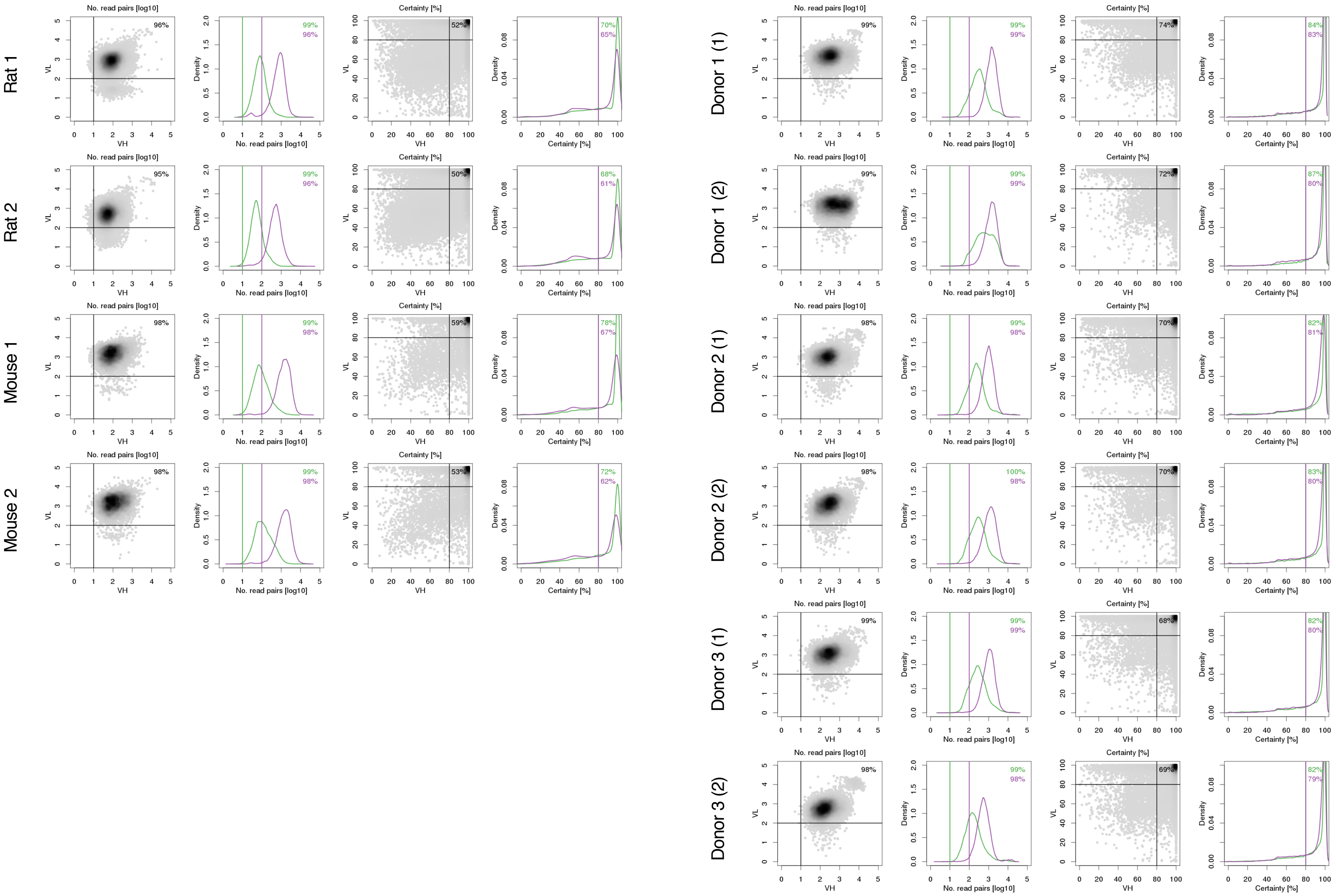

**Supplementary Figure 6**

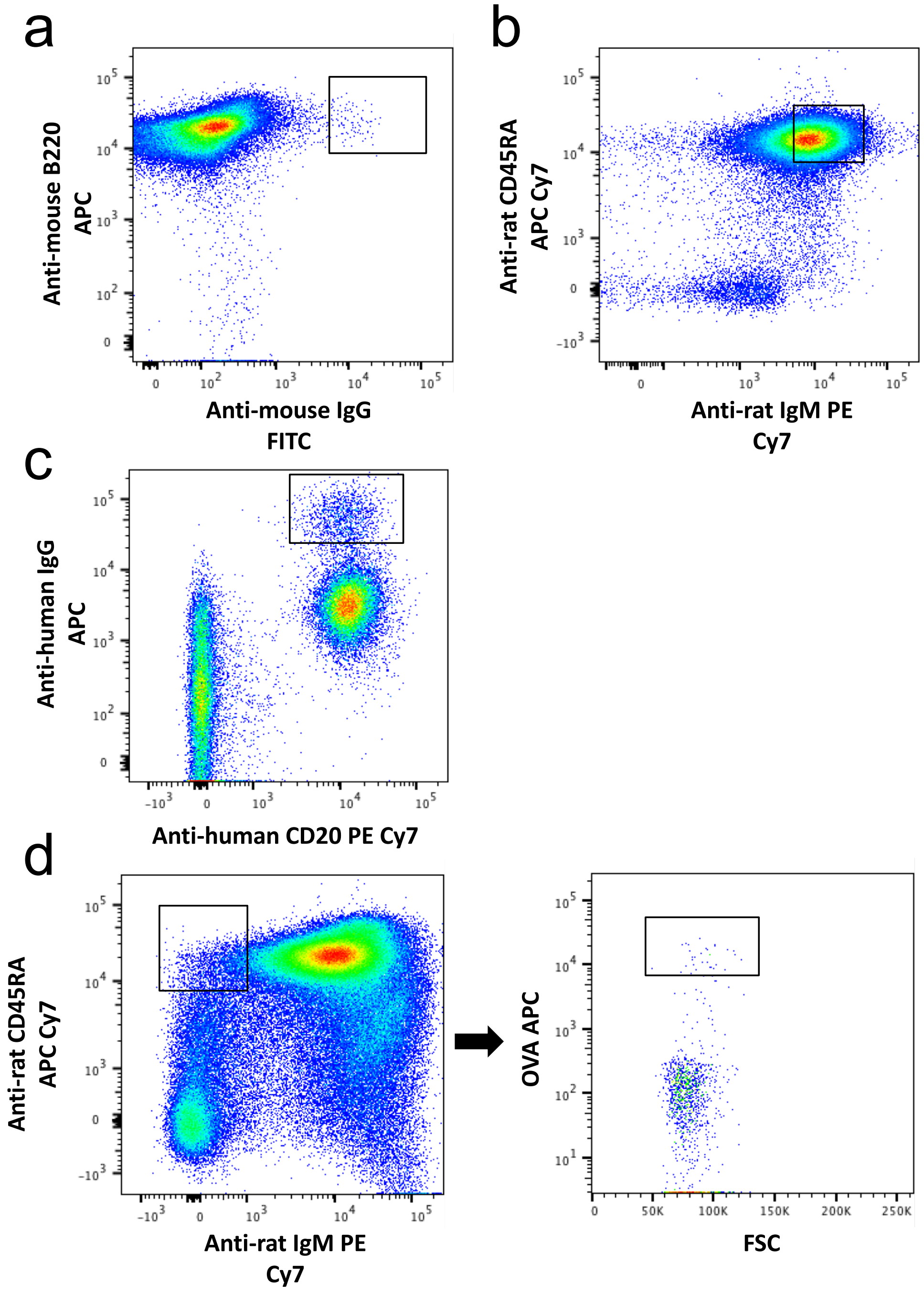

### Supplementary Figure 7

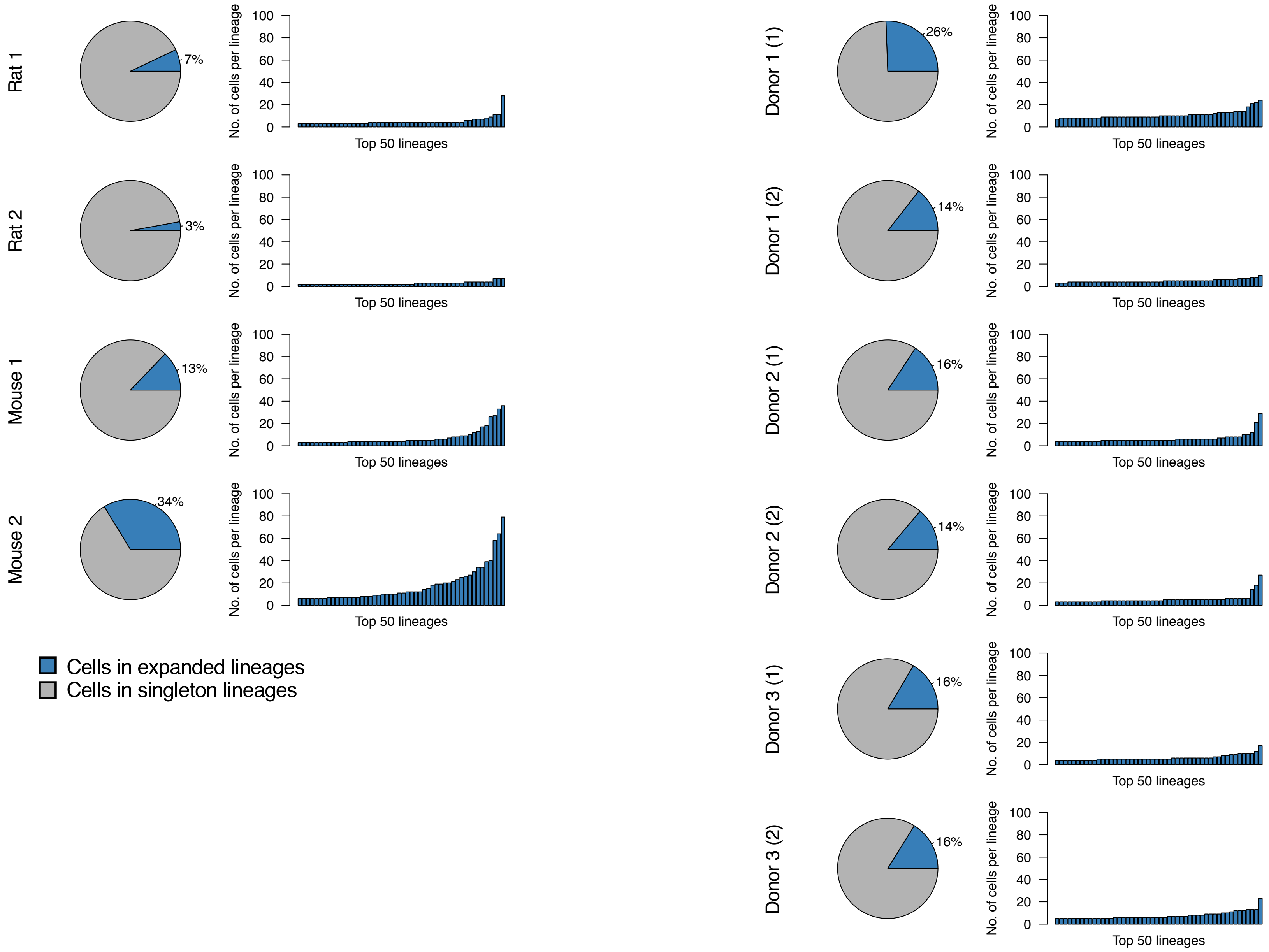

#### Supplementary Figure 8

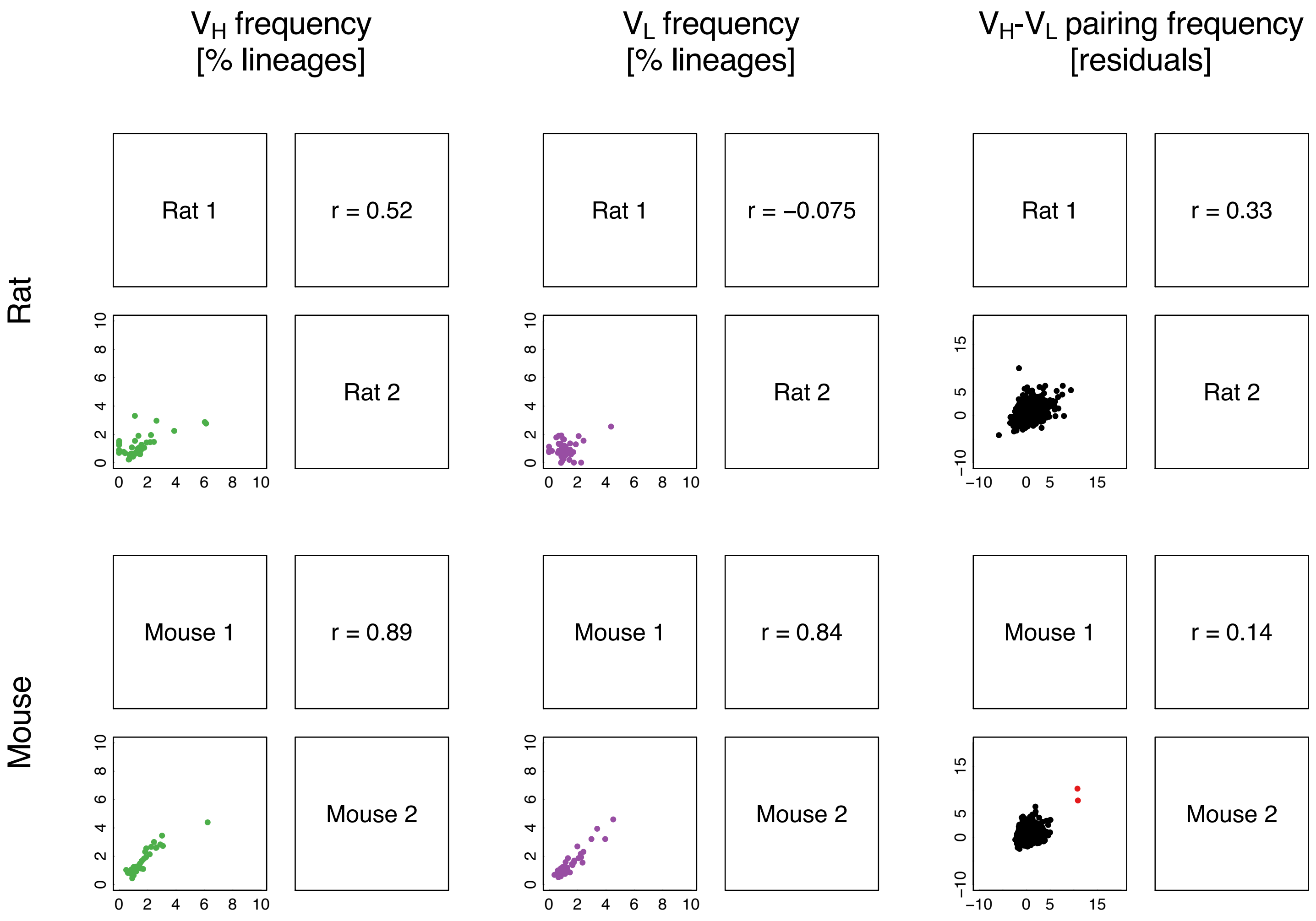

### Supplementary Figure 9

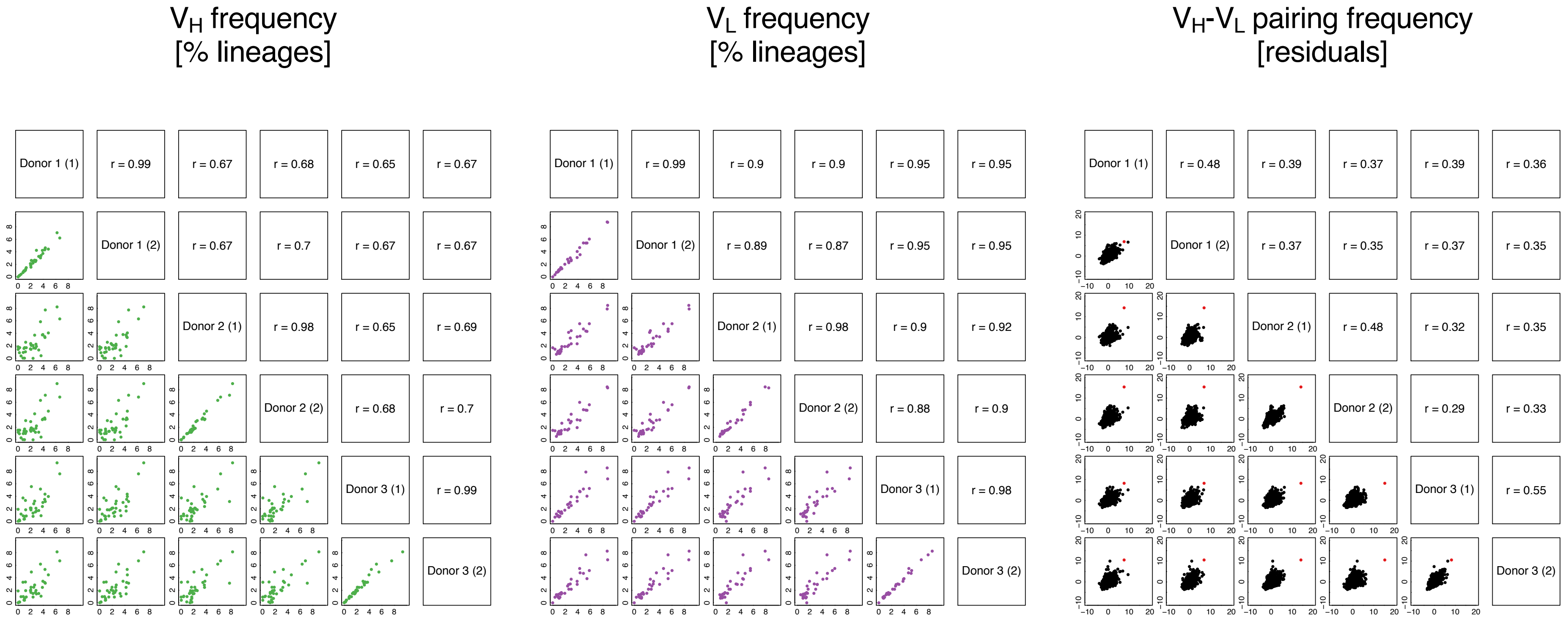

#### Supplementary Figure 10

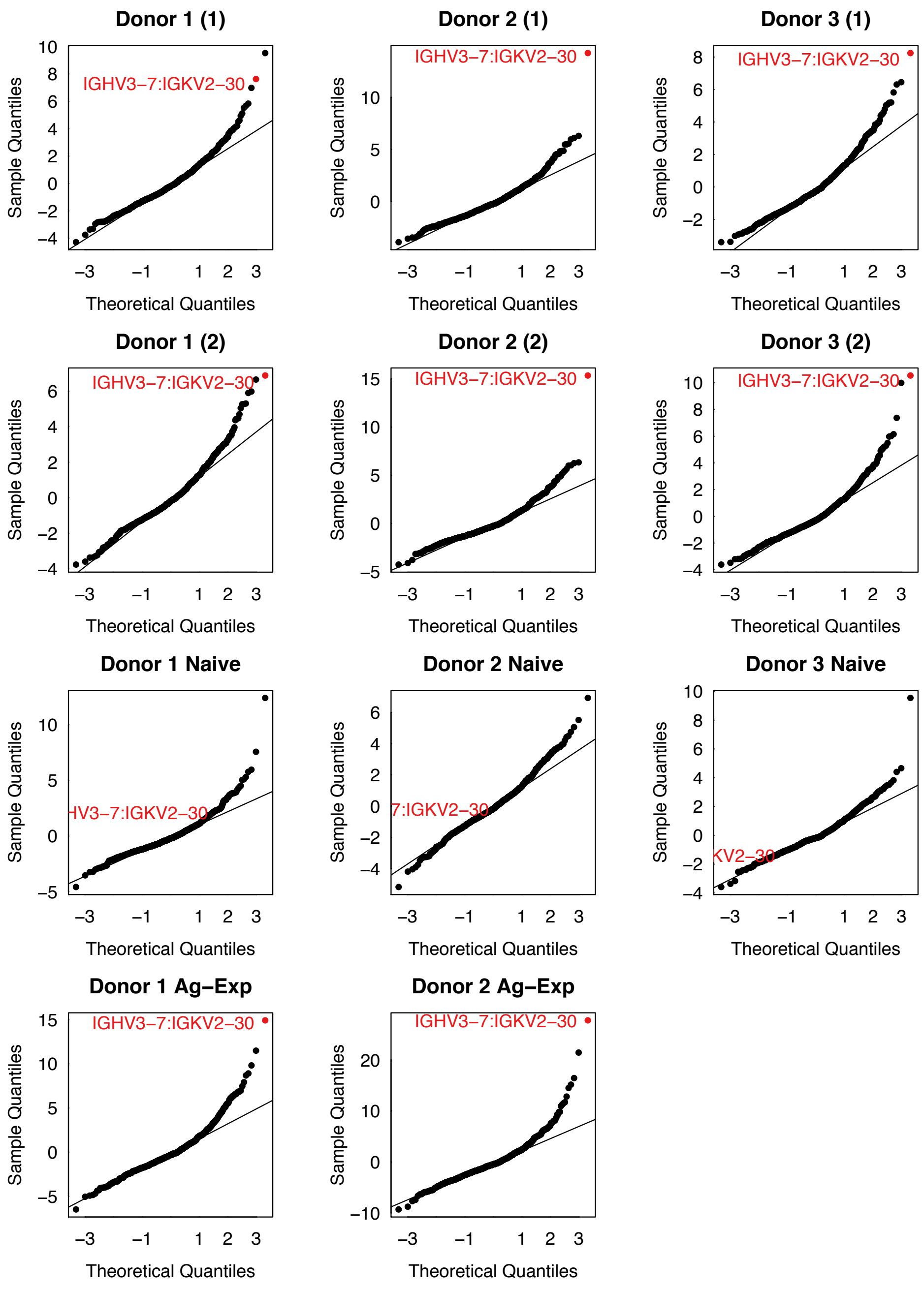

### Supplementary Figure 11

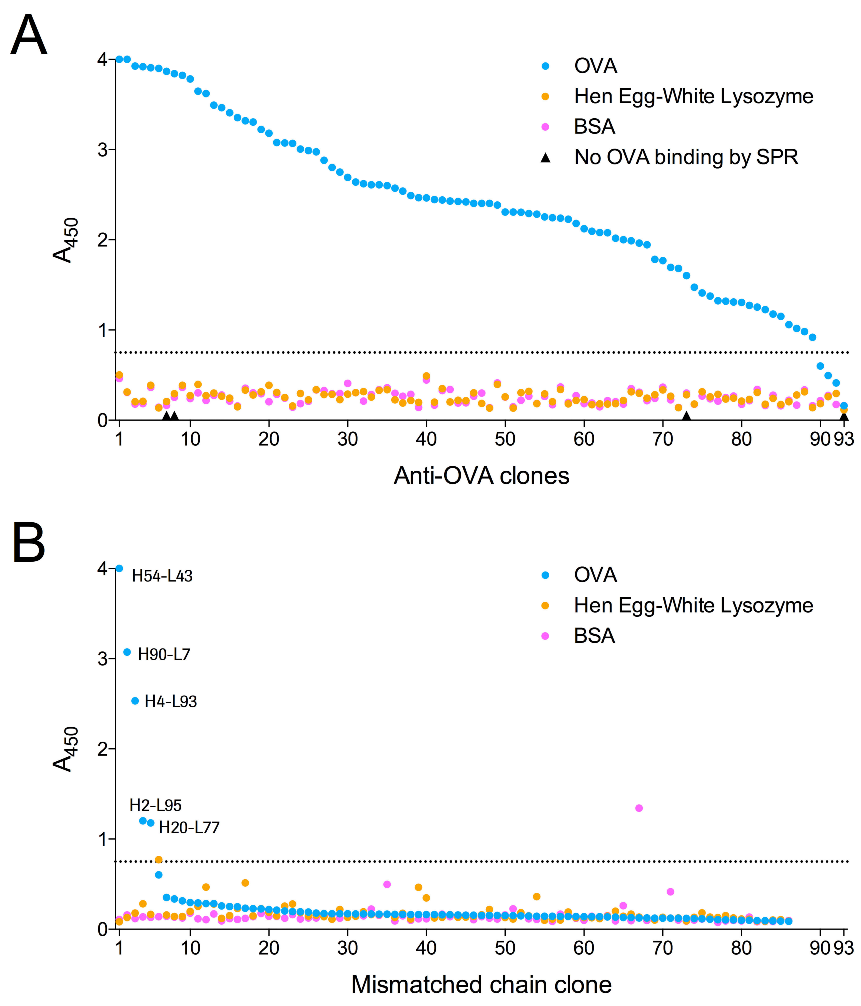

### Supplementary Figure 12

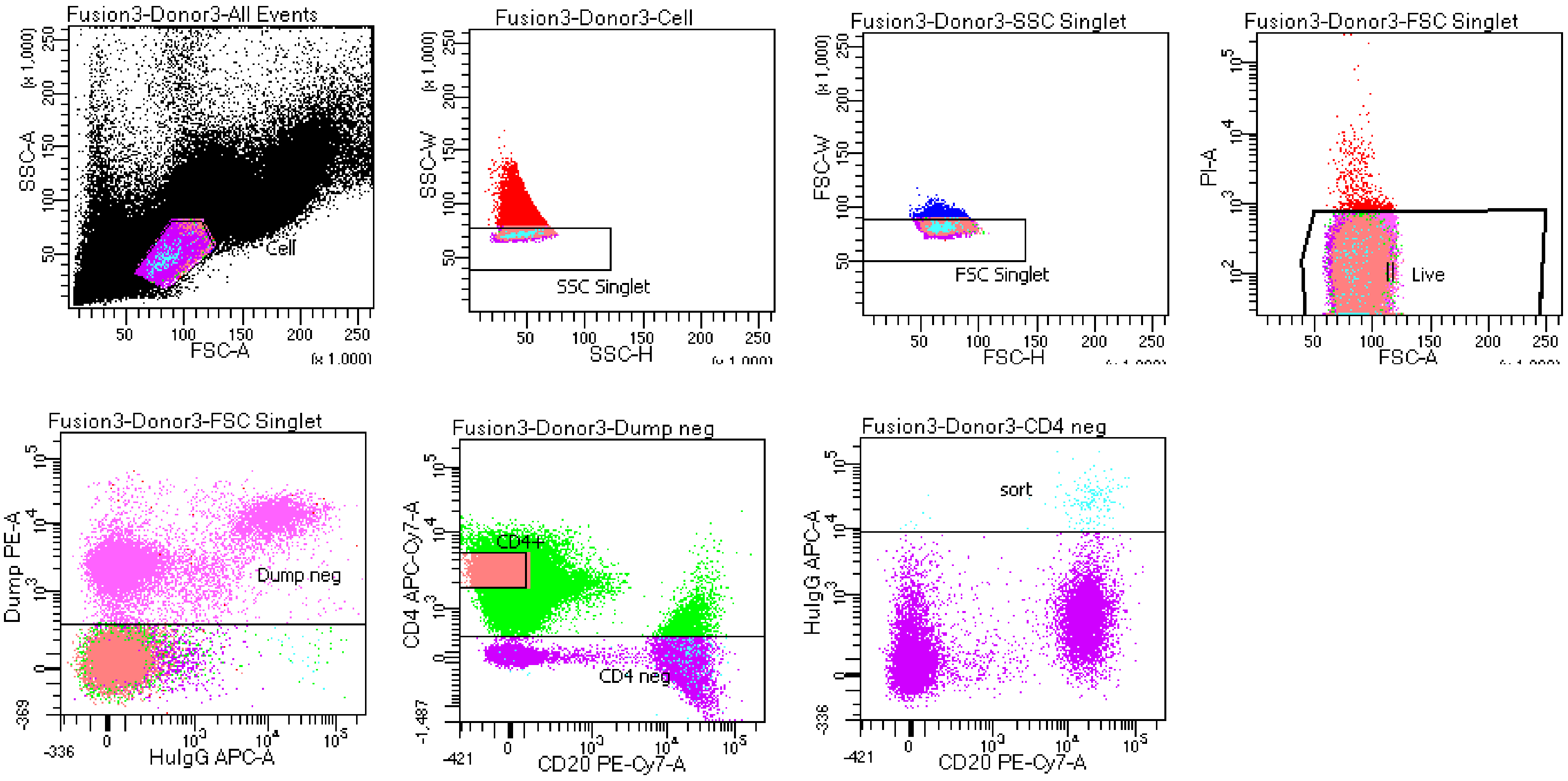


Tube: Donor3

| Population | #Events | %Parent | %Total |
|---|---|---|---|
| All Events | 510,250 | #### | 100.0 |
| Cell | 250,147 | 49.0 | 49.0 |
| SSC Singlet | 233,951 | 93.5 | 45.9 |
| FSC Singlet | 231,564 | 99.0 | 45.4 |
| Live | 230,891 | 99.7 | 45.3 |
| Dump neg | 113,173 | 49.0 | 22.2 |
| CD4 neg | 11,722 | 10.4 | 2.3 |
| sort | 178 | 1.5 | 0.0 |
| CD4+ | 49,323 | 43.6 | 9.7 |

**Supplementary Figure 13**

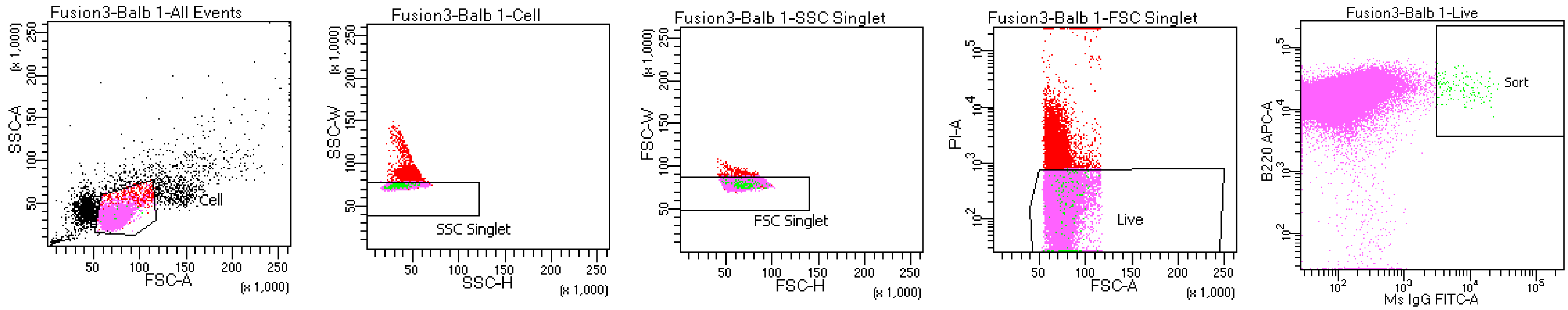


Tube: Balb 1

| Population | #Events | %Parent | %Total |
|---|---|---|---|
| All Events | 100,000 | #### | 100.0 |
| Cell | 78,224 | 78.2 | 78.2 |
| SSC Singlet | 74,841 | 95.7 | 74.8 |
| FSC Singlet | 74,547 | 99.6 | 74.5 |
| Live | 69,670 | 93.5 | 69.7 |
| Sort | 141 | 0.2 | 0.1 |

**Supplementary Figure 14**

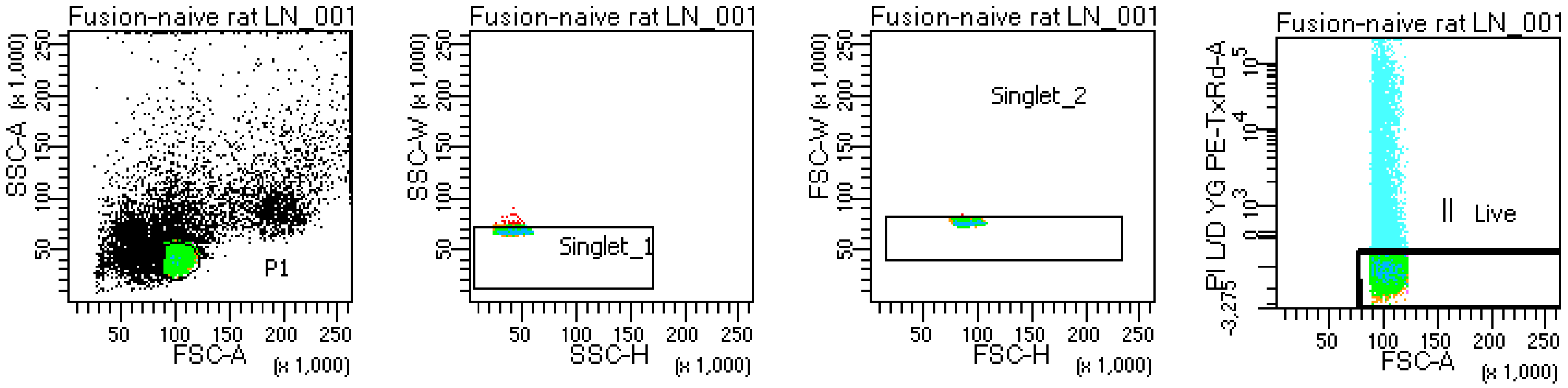


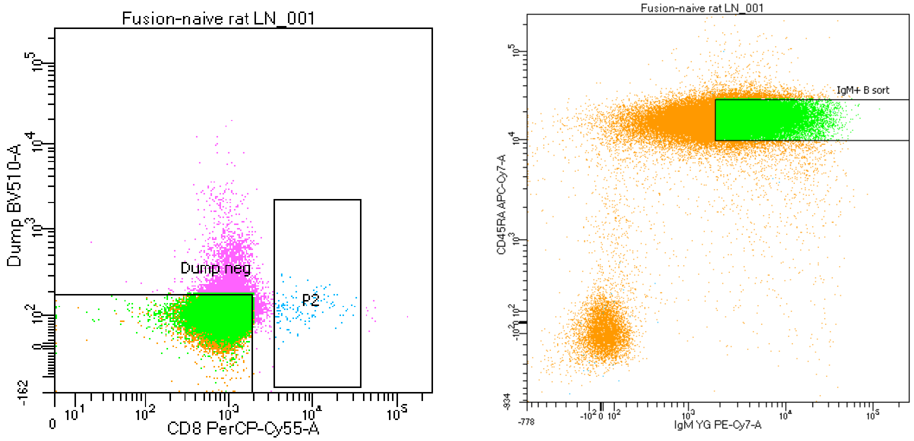


Tube: naive rat LN_001

| Population | #Events | %Parent | %Total |
|---|---|---|---|
| All Events | 243,300 | #### | 100.0 |
| P1 | 96,263 | 39.6 | 39.6 |
| Singlet_1 | 96,219 | 100.0 | 39.5 |
| Singlet_2 | 96,218 | 100.0 | 39.5 |
| Live | 87,696 | 91.1 | 36.0 |
| Dump neg | 83,716 | 95.5 | 34.4 |
| IgM+ B sort | 49,498 | 59.1 | 20.3 |
| P2 | 133 | 0.2 | 0.1 |

### Supplementary Figure 15

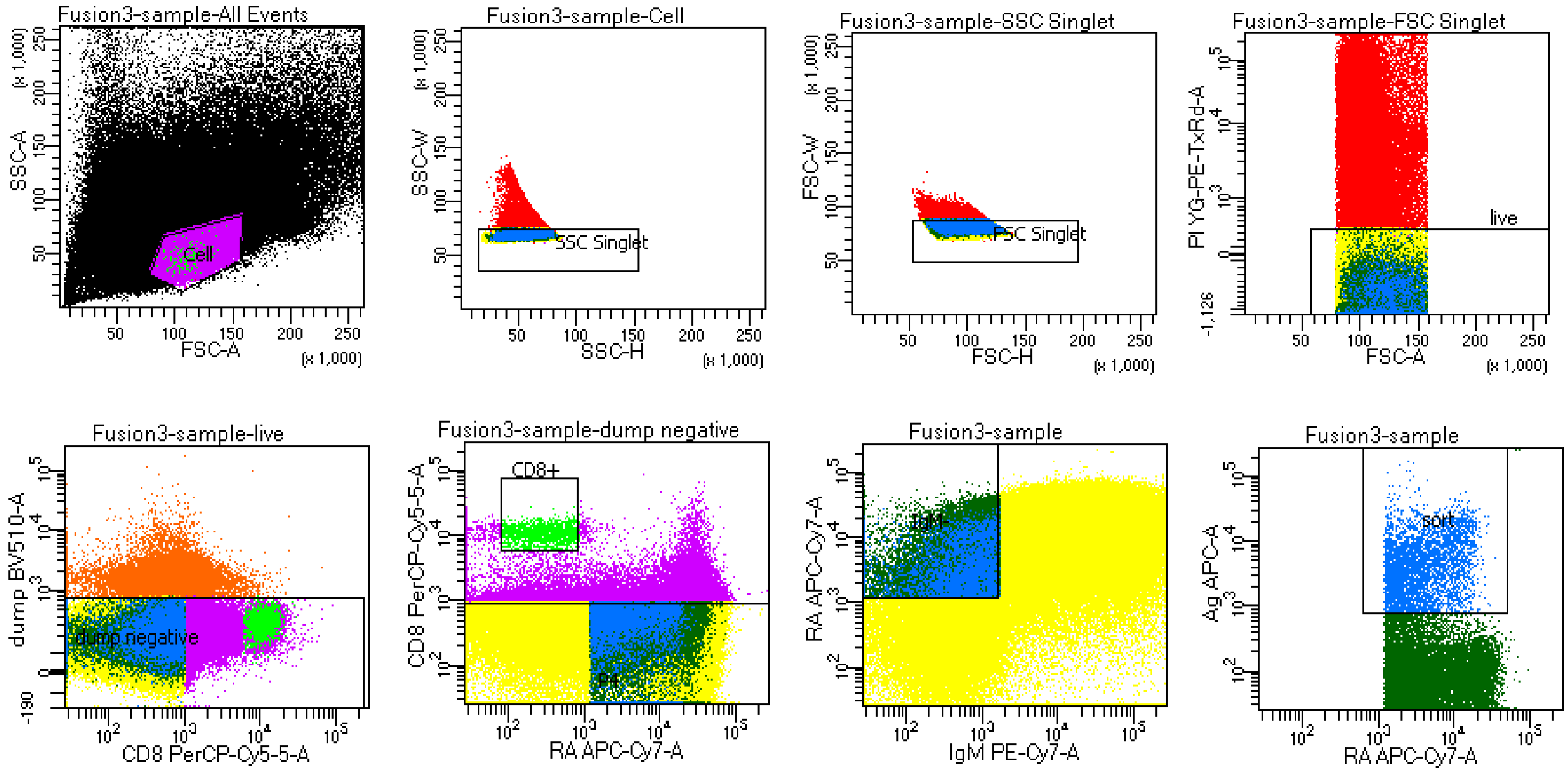


Tube: sample

| Population | #Events | %Parent | %Total |
|---|---|---|---|
| All Events | 4,225,000 | #### | 100.0 |
| Cell | 1,328,095 | 31.4 | 31.4 |
| SSC Singlet | 1,304,252 | 98.2 | 30.9 |
| FSC Singlet | 1,287,995 | 98.8 | 30.5 |
| live | 1,194,887 | 92.8 | 28.3 |
| dump negative | 1,141,436 | 95.5 | 27.0 |
| CD8+ | 1,391 | 0.1 | 0.0 |
| P4 | 1,013,025 | 88.8 | 24.0 |
| IgM- | 39,402 | 3.9 | 0.9 |
| sort | 4,356 | 11.1 | 0.1 |

### Supplementary Figure 16

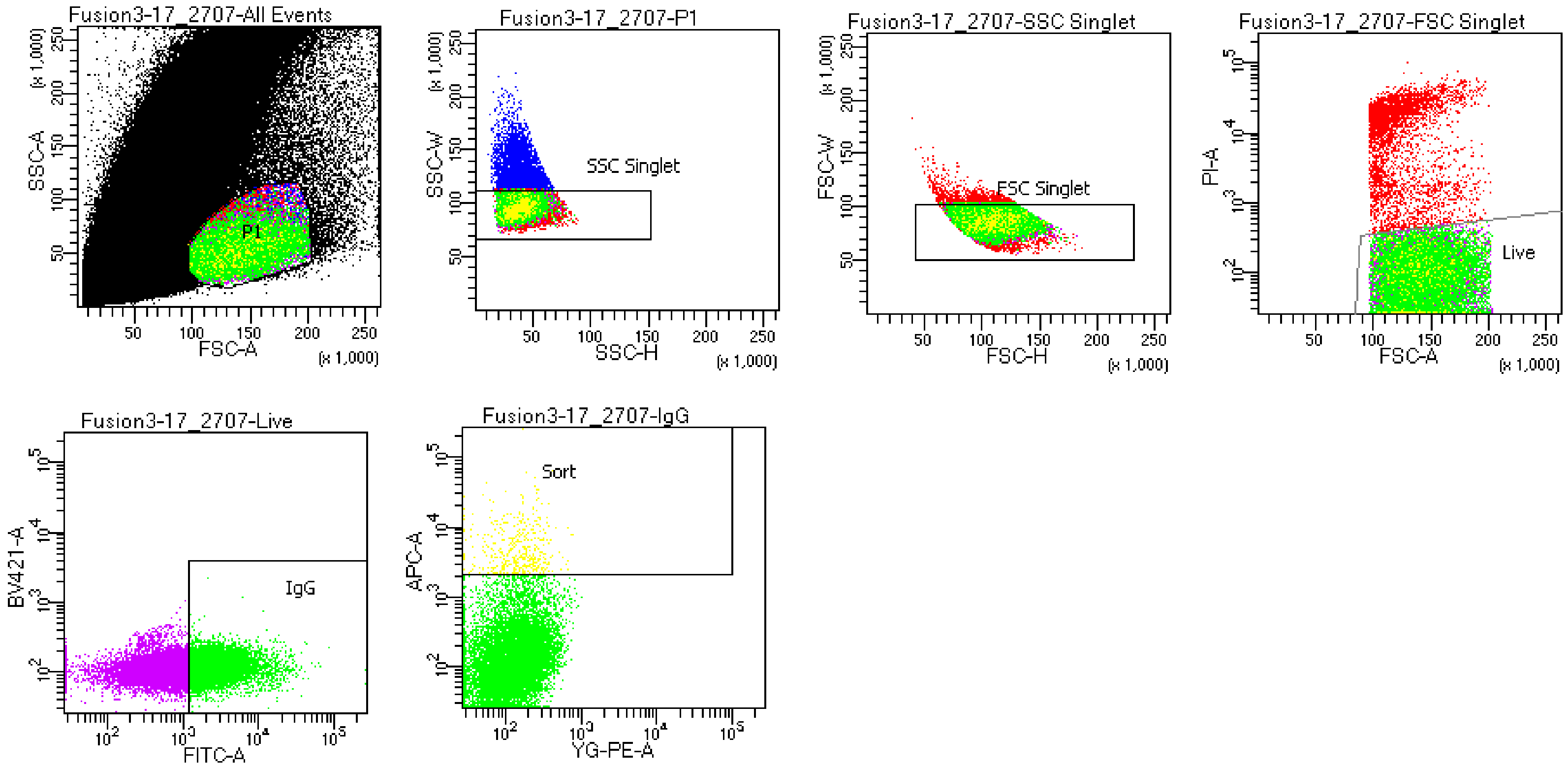


Tube: 17_2707

| Population | #Events | %Parent | %Total |
|---|---|---|---|
| All Events | 4,875,000 | #### | 100.0 |
| P1 | 36,297 | 0.7 | 0.7 |
| SSC Singlet | 32,718 | 90.1 | 0.7 |
| FSC Singlet | 31,846 | 97.3 | 0.7 |
| Live | 27,820 | 87.4 | 0.6 |
| IgG | 13,787 | 49.6 | 0.3 |
| Sort | 429 | 3.1 | 0.0 |
| NOT(P1) | 4,838,703 | 99.3 | 99.3 |
