## Supplementary Information for "Massively parallel single-cell B-cell receptor sequencing enables rapid discovery of diverse antigen-reactive antibodies"

**PCR primers (reverse strand, constant region)**

Rat CH1, outside PCR primer: YYTKGACCAGGCAKCCCAKDGTCAC
Rat CH1, inside PCR primer: CCAGGAGCCAGTGGATAGAC
Rat Cκ, outside PCR primer: AGGCACCTCCAGTTGCTAAC
Rat Cκ, inside PCR primer: CAGTTGGTGCAGCATCAGCCCG

Mouse CH1, outside PCR primer: GGGAARTAVYCCTTGACCAGGCABCC
Mouse CH1, inside PCR primer: GRCCARKGGATAGACHGATG
Mouse Cκ, outside PCR primer: GACTGAGGCACCTCCAGATG
Mouse Cκ, inside PCR primer: CTGCTCACTGGATGGTGGGAAG

**List of additional Rat VH germlines not in IMGT database**

>rIGHV2U2 ; wgs
CAGGTGCAGCTGAAGGAGTCAGGACCTGGTCTGGTGCAGCCCTCAGAGACCCTGTCCCTCACCTGCACT
GTCTCTGGGTTCTCATTAACCAGCAATAGTGTACACTGGGTTCGCCAGCCTCCAGGAAAGGGTCTGGAG
TGGATGGGAATAATACGGAGTGGTGGAAGCACAGATTATAATTCAGCTCTCAAATCCCGACTGAGCATC
AGCAGGGACACCTCCAAGAGCCAAGTTTTCTTAAAAATGAACAGTCTGCAAACTGAAGACACAGCCATT
TACTACTGTACCAGA

>rIGHV2U3 ; wgs
CAGGTGCAGCTGAAGGAGTCAGGACCTGGTCTGGTGCAGCCCTCACAGACCCTGTCCCTCACCTGCACT
GTCTCTGGATTCTCATTAACCAGCTATCATGTAAGCTGGGTTTGCCAGCCTCCAGGAAAGGGTCTGGAGT
GGATGGGAAGAATACAGAGTGGTGGAAGCACAGATTATAATTCAGCTCTCAAATCCCGACTGAGCATCA
GCAGGGACACCTCCAAGAGCCAAGTTTTCTTAAAAATGAACAGTCTGCAAACTGAAGACACAGCCATTT
ACTACTGTACCAGAGA

>rIGHV2U4 ; wgs
CAGGTGCAGCTGAAGGAGTCAGGACCTGGTCTGGTGCAGCCCTCACAGACTTTGTCTCTCACCTGCACT
GTCTCTGGGTTCTCACTAACCAGCTATCATGTAAGCTGGGTTCGCCAGCCTCCAGGAAAAGGTCTGGAG
TGGATGGGAGTAATATGGACTGGTGGAAGCACAGCATATAATTCACTTCTCAAATCCCGACTGAGCATC
AGCAGGGACATCTCCAAGAGCCAAGTTTTCTTAAAAATGAACAGTCTGCAAACTGAAGACACAGCCACT
TACTACTGTGCCAGAGA

>rIGHV2U5 ; wgs
CAGGTGCAGCTGAAGGAGTCAGGACCTGGTCTGGTGCAGCCCTCACAGACCCTGTCTCTCACCTGCACT
GTCTCTGGGTTCTCACTAACCAGCTATCATGTAAGCTGGGTTCGCCAGCCTCCAGGAAAGGGTCTGGAG
TGGATGGGAGGAATATGGGGTGATGGAAGCACAGCATATAATTCACTTCTCAAATCCCGACTGAGCATC
AGCAGGGACACCTCCAAGAGCCAAGTTTTCTTAAAAATGAACAGTCAGCAAACTGAAGACACAGCCATT
TACTTCTGTACCAGA

>rIGHV2U6 ; wgs

CAGGTGCAGCTGAAGGAGTCAGGACCTGGTCTGGTGCAGCCCTCACAGACCCTGTCCCTCACCTGCACT
GTCTCTGGGTTCTCACTAAGCAGCTATGGTGTAATCTGGGTTCACCAGCCTCCAGGAAAGGGTCTGGAG
TGGATGGGAGGAATATGGGGTGATGGAAGCACAGATTATAATTCAGCTCTCAAATCCCGACTGAGCATC
AGCAGGGACACCTCCAAGAGTCAAGTTTTCTTAAAAATGAACAGTCTGCAAACTGAAGACACAGCCATT
TACTTCTGTACCAGA

>rIGHV2U8 ; wgs
CAGGTGCAGCTGAAGGAGTCAGGATCTGGTCTGGTGCAGCCCTCACAGACCCTGTCCCTCACCTGCACT
GTCTCTGGGTTCTCACTAACCAGCTATGGTGTAAGCTGGGTTCGCCAGCCTCCAGAAAAGGGTCGGGAG
TGGATTGGAGCAATATGGACTGGTGGAAGCACAGATTATAATTCAGCTCTCAAATCCCGACTGAGCATC
AGCAGGGACACCTCCAAGAGCCAAGTTCTCTTAAAAATGAACAGTCTGCAAACTGAAGACACAGCCATG
TACTTCTGTGCCAGAAA

>rIGHV2U9 ; wgs
CAGGTACAACTGAAGGAGACAGGACCTGACCTAGTGCAACTGACACAGACCCTGTCCATCACATGCACT
GTCTCTGGGTTCTCATTAACCACCTATAATGTTCACTGGGTCCGTCAGCCTCCAGGAAAGGGTCTGGAGT
GGATGGGAGCAATGTGGAATGGTGGAGGCACAGATTATAATTCAGCATTTAAATCCCGACTGAGTATCA
GCAGGGACACCTCCAAGAGCCAAGTTTTCTTAAAAATGAACAGTTTGCAAACTGATGACACAGCCAAGT
ACTTCTGTGCCAGAA

>rIGHV2U12 ; wgs
CAGGTGCAGCTGAAGGAGTCAGGACCTGGTCTGGTGCAGCCCTCAGAGACCCTGTCCCTCACCTGCACT
GTCTCTGGGTTCTCACTAACCAGCTATAGTGTAAGTTGGGTTCGCCAGCCTTCAGGAAAAGGTCCTGAGT
GGATGGGAAGAATGTGGTATGATGGAGACACAGCATATAATTCAGCTCTCAAATCCCGACTGAGCATCA
GCAGGGACACCTCCAAGAACCAAGTTTTCTTAAAAATGAACAGTCTGCAAACTGATGACACAGGCACTT
ACTACTGTACCAGAGA

>rIGHV2U14 ; wgs
CAGGTGCAGCTGAAGGAGTCAGGACCTGGCCTGGTGAAGCCCTCAGAGACCCTGTCTCTCACCTGCACT
GTCTCTGGGTTCTCATTAACCAGCTATCATGTAAGCTGGGTTCGACAGCCTCCAGGAAAGGGTCTGGAG
TGGATGGGAGTAATATGGGGTGATGGAAGCACAGCATATAATTCAGCTCTCAAATCCCGACTGAGCATC
AGCAGGGACACCTCCAAGAGCCAAGTTTTCTTAAAAATGAGCAGTCTGAAAACTGAAGACACAGCCACT
TACTACTGTGCCAGAGA

>rIGHV2U16 ; wgs
CAGGTGCAGCTGAAGGAGTCAGGACCTGGTCTGGTGCAGCCCTCACAGACCCTGTCCCTCACCTGCACT
GTCTCTGGGTTCTCACTAACCAGCTATGGTGTAAGCTGGGTTCGCCAGCCTCCAGGAAAGGGTCTGGAG
TGGATGGGAGGAATATGGGGTGATGGAAGCACAAATTATAATTCAGCTCTCAAATCCCGACTGAGCATC
AGCAGGGACACCTCCAAGAGCCAAGTTTTCTTAAAAATGAACAGTCTGCAAACTGAAGACACAGCCATT
TACTTCTGTACCAGAGA

>rIGHV5U21 ; wgs
GAGGTGCAGCTGGTGGAGTCTGGGGGAGGCTTAGTGCAGCCTGGAAGGTCCATGAAACTCTCCTGTGC
AGCCTCAGGATTCACTTTCAGTAACTATGACATGGCCTGGGTCCGCCAGGCTCCAAAGAAGGGTCTGGA

GTGGGTCGCAACCATTAGTTATGATGGTAGTAGCACTTACTATCGAGACTCCGTGAAGGGCCGATTCAC
TATCTCCAGAGATAATGCAAAAAGCACCCTATACCTGCAAATGGACAGTCTGAGGTCTGAGGACACGGC
CACTTATTACTGTACAACAGA

>rIGHV5U22 ; wgs
GAGGTGCAGCTGGTGGAGTCTGGGGGAGGCTTAGTGCAGCCTGGAAGGTCCATGAAACTCTCCTGTGC
AGCCTCTGGATTCACTTTCAGTAGCTTTGCAATGGCCTGGGTCCGCCAGGCTCCAACGAAGGGGCTGGA
GTGGGTCGCATCCATTAGTTATGATGGTGGTAACACTTACTATCGAGACTCCGTGAAGGGCCGATTCACT
ATCTCCAGAGATAATGCAAAAAGCAGCCTATACCTGCAAATGGACAGTCTGAGGTCTGAGGACACGTCC
ACTTATTACTGTGCAAAAGACA

>rIGHV5U23 ; wgs
GAGGTGCAGCTGGTGGAGTCTGGGGGCGGCTTAGTGCAGCCTGGAAGGTCCATGAAACTCTCCTGTGC
AGCCTCAGGATTCACTTTCAGTAACTATGGCATGGCCTGGGTCCGCCAGGCTCCAAAGAAGGGTCTGGA
GTGGGTCGCATACATTAGTTATGATGGTGGTAGCACTTACTATCGAGACCCCGTGAAGGGCCGATTCAC
TATCTCCAGAGATAATGCAAAAAGCACCCTATACCTGCAAATGGACAGTCTGAGGTCTGAGGACACGGC
CACTTATTACTGTACAACAGA

>rIGHV5U25 ; wgs
GAGGTGCAGCTGGTGGAGTCTGGGGGAGGATTAGTGCAGCCTGGAAGGTCCCTGAAACTCTCCTGCGC
AGCCTCAGGATTCACTTTCAGTAGCTTTCCAATGGCCTGGGTCCGCCAGGCTCCAAAGAAGGGTCTGGA
GTGGGTCGCATCCATTAGTTCTGGTGGTGGTGGCACTTACTATCCAGATTCCGTGAAGGGCAGATTTACT
ATCTCCAGAGATAATGCAAAAAGCACCCTATACCTGCAAATGGACAGTCTGAGATCTGAGGACACGGCC
AGTTACTATTGTGCAAGACA

>rIGHV2U17d ; repertoire NGS
CAGGTGCAGCTGAAGGAGTCAGGACCTGGTCTGGTGCAGCCCTCACAGACCCTGTCCCTCACCTGCACT
GTCTCTGGGTTCTCATTAACCAGCAATAGTGTACACTGGGTTCGCCAGCCTCCAGGAAAGGGTCTGGAG
TGGGTTGGAGCAATATGGAGTGGTGGAAGCACAGATTATAATTCAGCTCTCAAATCCCGACTGAGCATC
AGCAGGGACACCTCCAAGAGCCAAGTTTTCTTAAAAATGAACAGTCTGCAAACTGAAGACACAGCCATT
TACTTCTGTACCAGA

>rIGHV2U18d ; repertoire NGS
CAGGTGCAGCTGAAGGAGTCAGGACCTGGTCTGGTGCAGCCCTCACGGACCCTGTCCCTCACCTGCACT
GTCTCTGGGTTCTCGCTCACCAGCTATGGTGTAAGCTGGGTTCGCCAGCCTCCAGGAAAGGGTCTGGAG
TGGATTGCAGCAATATGGAGTGGTGGAAGTACAGATTATAATTCAGCTCTCAAATCCCGACTAAGCATC
AGCAGGGACACCTCCAAGAGCCAAGTTCTCTTAAAAATGAACAGTCTGCAAACTGAAGACACAGCCATG
TACTTCTGTGCCAGA

>rIGHV3U1d ; repertoire NGS
GAGGTGCAGCTTCAGGAGTCAGGACCTGGCCTTGTGAAACCCTCACAGTCACTCTCCCTCACCTGTTCTG
TCACTGGTTACTCCATCACTAGTAATTACTGGGGCTGGATCCGGAAGTTCCCAGGAAATAAAATGGAGT
GGATGGGATACATAAGCTACAGTGGTAGCACTAGCTACAACCCATCTCTCAAAAGTCGAATCTCCATTAC

TAGAGACACATCGAAGAATCAGTTCTTCCTGCAGTTGAACTCTGTAACTACTGAGGACACAGCCACATAT
TACTGTGCAAGA

>rIGHV3U2d ; repertoire NGS
GAGATACAGCTGCAGGAGTCAGGACCTGGCCTTGTGAAACCTTCACAGTCACTCTCCCTCACCTGTTCTG
TCACTGGTTACACCATTACCAGTGGTTATGATTGGAGCTGGATCCGGAAGTTCCCAGGAAATAAAATGG
AGTGGATGGGATACATAAGCTACAGTGGTAGCACTAACTACAACCCATCGCTCAAAAGTCGAATCTCCA
TTACTAGAGACACATCCAAGAATCAGTTCTTCCTGCAGTTGAACTCTGTAACTACTGAGGATACAGCCAC
ATATTACTGTGCAAGA

>rIGHV5U27d ; repertoire NGS
GAGGTGCAGCTGGTGGAGTCTGGCGGAGGCTTAGTACAGCCTGGAAGGTCCCTGAAACTCTCCTGTGC
AGCCTCAGGATTCACTTTCAGTAACTATTACATGGCCTGGGTCCGCCAGGCTCCAACGAAGGGTCTGGA
GTGGGTCGCATCCATTACTAATAGTGGTGGTAGTACTTACTATCGAGACTCCGTGAAGGGCCGATTCACT
ATCTCCAGAGATAATGCAAAAAGCACCCTATACCTGCAAATGGACAGTCTGAGGTCTGAGGACACGGCC
ACTTATTACTGTACAAGA

>rIGHV5U28d ; repertoire NGS
GAGGTGCAGCTGGTGGAGTCTGGGGGCGGCTTGGTGCAGCCTGGAAGGTCCCTGAAACTCTCCTGTGC
AGCCTCAGGATTCACTTTCAGTAACTATGACATGGCCTGGGTCCGCCAGGCTCCAACGAAGGGTCTGGA
GTGGGTCGCATCCATTAGTCCTAGTGGTGGTAGCACTTACTATCGAGACTCCGTGAAGGGCCGATTCAC
TATCTCCAGAGATAATGCAAAAAGCACCCTATACCTGCAAATGGACAGTCTGAGATCTGAGGACACCGC
CACTTATTACTGTGCAACG

>rIGHV5U29d ; repertoire NGS
GAGGTGCAGCTGGTGGAGTCTGGGGGAGGCTTAGTGCAGCCTGGAAGGTCCCTGAAACTCTCCTGTGC
AGCCTCAGGATTCACTTTCAGTGACTATAACATGGCCTGGGTCCGCCAGGCTCCAACGAAGGGTCTGGA
GTGGGTTGCATCCATTAGTTATGGTGGTAGTAACACCCACTATGGAGACTCCGTGAAGGGCCGATTCAC
TATCTCCAGAGATAATGCAAAAAGCACCCTATACCTGCAAATGGACAGTCTGAGGTCTGAGGACACGGC
CACTTATTACTGTGCAAGA

>rIGHV5U30d ; repertoire NGS
GAGGTGCAGCTGGTGGAGTCTGGGGGAGGCCTAGTGCAGCCTGGAAGGTCCCTGAAACTCTCCTGTGC
AGCCTCAGGATTCACTTTCAGTAACTATTACATGGCCTGGGTCCGCCAGGCTCCAAAGAAGGGTCTGGA
GTGGGTCGCAACCATTAGTACCAGTGGTAGCAGAACTTACTATCCAGACTCCGTGAAAGGCCGATTCAC
TATCTCCAGAGATAATGCAAAAAGCAGCCTATACCTGCAAATGAACAGTCTGAAGTCTGAGGACACGGC
CACTTATTACTGTGCAAGA

>rIGHV5U31d ; repertoire NGS
GAGGTGCAGCTGGTGGAGTCTGGGGGAGGCTTAGTGCAGCCTGGAAGGTCCATGAAACTCTCCTGTGC
AGCCTCAGGATTCACTTTCAATAACTATGACATGGCCTGGGTCTGCCAGGCTCCAAAGAAGGGTCTGGA
GTGGGTCGCAACCATTAGTTATGATGGTAGTAGCACTTACTATCGAGACTCCGTGAAGGGCCGATTCAC
TATCTCCAGAGATAATGCAAAAAGCACCCTATACCTGCAAATGGACAGTCTGAGGTCTGAGGACACGGC
CACTTATTACTGTGCAAGA

**List of additional Rat Vκ germlines not in IMGT database**

>rIGKV1U5 ; wgs
GATGTTGTGATGACCCAGACACCACCATCTTTGTCGGTTGCCATTGGACAGTCAGTCTCCATCTCTTGCA
AGTCAAGTCAAAGCCTCGTAGCTAGTGATAAAAATACATATTTGAATTGGTTATTACAGAGTCCTGGCCG
GTCTCCGAAGCGCCTAATCTATCAGGTGTCTAAGCTGGACTCTGGAGTCCCTGACAGGTTCAGTGGCAG
TGGATCAGAGAAAGATTTCACACTTAAAATCAGCAGAGTGGAGGCTGAGGATTTGGGAGTTTATTACTG
CCTGCAAGGTACACATCTTCCTCA

>rIGKV1U10 ; wgs
GATGTTGTGTTGACACAAACTCCAGTTGCCCAGCCTGTCACACTTGGAGATCAAGCTTCTATATCTTGCA
GGTCTAGTCAGAGCCTGGTACATAGTAATGGAAACACTTATTTGGAATGGTACCTACAGAAGCCAGGCC
AGTCTCCACAGCTCCTCATCTATAAGGTTTCCAACCGATTTTCTGGGGTACCAGACAGGTTCATTGGCAG
TGGGTCAGGGTCAGATTTCACCCTCAAGATCAGCAGAGTAGAGCCTGAGGACTTGGGAGTTTATTACTG
CTTCCAAGCTACACATGATCCTCC

>rIGKV1U75 ; wgs
GATGTTGTGTTGACACAAACTCCAGGTTCCCTGTCTGTCACACTTGGAGATCAAGCTTCTATATCTTGCAG
GTCTAGTCAGAGCCTGGAATATAGTGATGGATACACTTATTTGGAATGGTACCTACAGAAGCCAGGCCA
GTCTCCACAGCTCCTCATCTATGAAGTTTCCAACCGATTTTCTGGGGTCCCAGACAGGTTCATTGGCAGT
GGGTCAGGGACAGATTTCACCCTCAAGATCAGCAGAGTAGAGCCTGAGGACTTGGGAGTTTATTACTGC
TTCCAAGCTACACATGATCCTCC

>rIGKV2U3 ; wgs
GATGTTGTGCTGACCCAGACTCCACCCACTTTATCGGCTACCATTGGACAATCGGTCTCCATCTCTTGCAG
GTCAAGTCAGAGTCTCTTAGATAGTGATGGAGATACCTATTTAAATTGGTTGCTACAGAGGCCAGGCCA
ATCTCCACAGCTTCTAATTTATTCGGTATCCAACCTGGAATCTGGGGTCCCCAACAGGTTCAGTGGCAGT
GGGTCAGAAACAGATTTCACACTCAAAATCAGTGGAGTGGAGGCTGAAGATTTGGGAGTTTATTACTGC
ATGCAAGCTACCCATGCTCCTCT

>rIGKV2U13 ; wgs
GATATTGTGATGACTCAAGCTCCACTCTCTGTATCTGTCACTCCTGGAGAGTCAGCTTCCATCTCCTGCAG
ATCTAGTAAGAGTCTGCTAAGTAGTAAGGGCATCACTTCCTTGTATTGGTACCTTCAGAGGCCAGGAAA
GTCTCCTCAGCTCCTGATATATCGGATGTCCAACCTTGCCTCAGGAGTTCCAGACAGGTTTAGTGGCAGT
GGGTCAGAAACCGATTTTACACTGAAAATCAGTAAGGTGGAGACTGAGGATGTTGGTGTTTATTACTGT
GGACATCGGTCTAGAATATCCTCC

>rIGKV2U17 ; wgs
GATATTGTGATGACCCAGGGTGCACTCCCCAATCCTGTCCCTTCTGGAGAGTCAGCTTCCATCACCTGCC
AGTCTAGTAAGAGTCTGCTGCACAGCAATGGCAAGACATACTTGAATTGGTATCTGCAGAGGCCAGGAC
AGTCTCCTCAGCTCCTGATCTATTGGATGTCTACCCGTGCATCAGGAGTCTCAGACAGGTTCAGTGGCAG
TGGGTCAGGAACAGATTTCACACTGAAAATCAGTAGCGTGGAGGCTGAGGATGTGGGTGTGTATTACT
GTCAGCAATTTCTAGAGTATCCTC

>rIGKV2U18 ; wgs
GATATTGTGTTGACTCAAGCTCCACTCTCTGTATCTGTCACTCCTGGAGAGTCAGCTTCCATCTCCTGCAG
GTCTAGTAAGAGTCTGCTAAGTAGTAAGGGCATCACTTCCTTGTATTGGTACCTTCAGAGGCCAGGAAA
GTCTCCTCAGCTCCTGATATATCAGATGTCCAACCTTGCCTCAGGAGTTCCAGACAGGTTTAGTAGCAGT
GGGTCAGAAACAGATTTTACACTGAAAATCAGTAAGGTGGAGACTGAGGATGTTGGTGTTTATTACTGT
GGACATCGTCTAGAATATCCTCC

>rIGKV3U19 ; wgs
GACATTGTGCTGACCCAGTCTCCTGCTTTGGCTGTGTCTCTAGAGCAGAGAGCCACCATCTCTTGCAAAA
CCAGCCAGAATGTCGATAATTATGGCATTAGTTATATGCACTGGTACCAACAGAAACCAGGACAGCAAC
CCAAACTCCTCATCTATGGTGCATCCAACCTAGAGTCTGGAGTCCCTGCCAGGTTCAGTGGCAGTGGGTC
TGGGACAGACTTCACCCTCACCATCGATCCTGTGGAGGCTGATGATATTGCAACCTATTACTGTCAGCAG
AGTAAGGATTATCCTCC

>rIGKV4U34 ; wgs
GAAATTGTGCTAACCCAGTCTCCAACAACCATGGCTGCATCTCCGGGGGAGAAGGTCACCATCACCTGC
CGTGCCAGCTCCAGTGTAAGCTACATGTACTGGTACCAGCAGAAGTCAGGCGCCTCCCCTAAACCCTGG
ATTTATGAAACATCCAAACTGGCTTCTGGAGTCCCAGATCGCTTCAGTGGCAGTGGGTCTGGGACCTCTT
ATTCGTTCACAATCAGCTCCATGGAGACTGAAGATGCTGCCACTTATTACTGTCACCAGTGGAGTAGTAC
CCCACCCA

>rIGKV8U28 ; wgs
GACATTGTGATGACCCAGACTCCATCCTCCCAGGCTGTGTCAGCAGGGGAGAAGGTCACTATGAGCTGC
AAGTCCAGTCAGAGTCTTTTATACAGTGGAGACCAAAAGAACTACTTGGCCTGGTACCAGCAGAAACCT
GGGCAGTCTCCTAAACTGCTGATCTACTTGGCATCCACTAGGGAATCAGGGGTCCCTGATCGCTTCATAG
GCAGTGGATCTGGGACAGACTTCACTCTGACCATCAGCAGTGTGCAGGCTGAAGATCTGGCAGATTATT
ACTGTCAGCAGCATTACAGCTATCCTCC

>rIGKV8U29 ; wgs
GACATTGTGATGACCCAGTCTCCCTCCTCCCTGGCTGTGTCAGCAGGAGACAAGGTCACTATGAGCTGC
AAGTCCAGTCAGAGTCTTTTATCCAGTGAATACCAAGGGAACTACTTGTCCTGGTTCCAGCAGAAACCAG
GGCAGTCTCCTAAACTGCTGATCTCCTTGGCATCCACTAGGGAAACTGGTGTCCCTGATCGCTTCATAGG
CAGTGGATCTGGGACAGACTTCACTCTGACCATCAGCAGTGTGCAGGCTGAAGACCTGGCTGTTTATTA
CTGTGAGCAGTACTACAGCTATCCTCC

>rIGKV8U40 ; wgs
ACATTGTGATGACCCAGACTCCATCCTCCCAGGCTGTGTCAGCAGGGGAGAAGGTCACTATGAGCTGCA
AGTCCAGTCAGAGTCTTTTATACAGTGAAAACAAAAAGAACTACTTGGCCTGGTACCAGCAGAAACCAG
GGCAGTCTCCTAAACTGCTGATCTACTGGGCATCCACTAGGGAATCTGGGGTCCCTGATCGCTTCATAGG
CAGTGGATCTGGGACAGATTTCACTCTGACCATCAGCAGTGTGCAGGCAGAAGACCTGGCTGTTTATTA
CTGCCAGCAGTACTATAACTTTCCTCC

>rIGKV10U55 ; wgs

GACATCAAGATGACCCAGTCTCCCTCCTCCCTGTCTGCATCTCTGGGAGAAAGAGTCACCATCAGTTGCA
GGGCAAGTGAGAATATTAACAATATTTGGCCTGGTATCAGAAGAAAGAAGATGGAAGTGTTAAACTCC
TGATTTACTACACATCAAATCTACAATCTGGGGTCCCATCAAGGTTCAGTGGCAGTGGGTCTGGGAAAG
ATTACTCTCTTACCATTAGTGGCCTAGAATCTGAAGATATTGCGACTTACTATTGTCAGCAAGGTTATACC
CCACC

>rIGKV12U30 ; wgs
GACATCCAGATGACACAGTCTCCAGCTTCCCTGTCTGCATCTCTGGGAGAAACTGTCACCATCGAATGTC
GAGCAAGTGAGGACATTTACAGTAATTTAGCGTGGTATCAGCAGAAACCAGGGAACTCTCCTCAGCTCC
TGATCTATGATGCAAATAGCTTGGCAGATGGGGTCCCATCACGGTTCAGTGGCAGTGGATCTGGCACAC
AGTATTCTCTAAAGATAAACAGCCTGCAATCTGAAGATGTCGCAAGTTATTTCTGTCAACAGTATAACAA
TTATCCTCC

>rIGKV12U74 ; wgs
GACATCCAGATGACACAGTCTCCAGCTTCCCTGTCTGCATCTCTGGGAGAAACTATCTCCATCGAATGTC
TAGCAAGTGAGGGCATTTCCAGTTATTTAGCGTGGTATCAGCAGAAGCCAGGGAAATCTCCTCAGCTCC
TGATCTATGGTGCAAATAGCTTGCAAGCTGGGGTCCCATCACGGTTCAGTGGCAGTGGATCTGGCACAC
AGTATTCTCTCAAGATCAGCAGCATGCAACCTGAAGATGAAGGGGATTATTTCTGTCAACAGAGTTACA
AGTTTCCTCC

>rIGKV12U76 ; wgs
GACATCCAGATGACACAGTCTCCTGCCTCCCTGTCTGCATCTCTGGAAGAAATTGTCACCATCACATGTC
AGGCAAGCCAGGACATTGGTAATTGGTTGGCATGGTATAAGCAGAAACCAGGGAAATCTCTTCAGCTCC
TGATCAATGATGCAACCAGCTTGGCAGACGGGGTCCCATCAAAGTTCAGAGGCAGTAGATCTGGTACAC
AGTATTCTCTTAAGATCAGCAGACTACAGGATGAAGATTTTGGAAGCTATTACTGTCAACAGGCTCATAG
TAATCCTCC

>rIGKV12U78 ; wgs
GACATCCAGATGACACAGTCTCCTGCCTCCCTGTCTGCTTCTCTGGAAGAAATTGTCACCATCACCTGCAA
GGCAAGCCAGGGCATTGATGATTACTTATCATGGTATCAGCAGAAACCAGGGAAATCTCCTCAGCTCCT
GATCTATGATGCAACCAGCTTGGCAGATGGGGTCCCATCACGGTTCAGCGGCAGTAGATCTGGCACACA
GTATTCTCTTAAGATCAGCAGACCACAGGTTGATGATTCTGGAATCTATTACTGTCTACAGAGTTACAGT
ACTCCTCC

>rIGKV12U81 ; wgs
GACATCCAGATGACACAGTCTCCTGCCTCCCTGTCTGCATCTCTGGAAGAAATTGTCACCATCACATTCCA
GGCAAGCCAAGACATTGGTAATTGGTTGGCATGGTATCAGCAGAAACCGGGGAAATCTCCTCAGCTCCT
GATTTATGATGCAACCAGCTTGGCAGATGGGGTCCCATCACGGTTCAGCGGCAGTAGATCTGGCACACA
GTATTCTCTTAAGATCAGCAGACTACAGGTTGAAGATATTGGAAGCTATTACTGTCAACAGGCTCATAGT
AATCCTCC

>rIGKV15U51 ; wgs
GACATCCAGATGACCCAGTCTCCTTCACTCCTGTCTGCATCTGTGGGAGACAGAGTCACTCTCAACTGCA
AAGCAAGTCAGAATATTTATAAGAACTTAGAGTGGTATCAGCAAAAGCATGGAGAAGCTCCAAAACTCC

TGATATATTATACAAACAATTTGCAAACGGGCATCTCATCAAGGTTCAGTGGCAGTGGATCTGGTACAG
ATTACACACTCACCATCAGCAGCCTGCAGCCTGAAGATGTTGCCACATATTACTGCTATCAGTATAACAG
CGGGCCC

>rIGKV16U47 ; wgs
GATGTCCAGATGACCCAGTCTCCGTCTTATCTTGCTGCGTCTCCTGGAGAAAGTGTTTCCATCAGTTGCA
AGGCAAGTAAGAGCATTAGCAATTATTTAGCCTGGTATCAACAGAAACCTGGGGAAGCAAATAAGCTTC
TTATCTACTCTGGGTCAACTTTGCAATCTGGAACTCCATCGAGGTTCAGTGGCAGTGGATCTGGTACAGA
TTTCACTCTCACCATCAGAAACCTGGAGCCTGAAGATTTTGCAGTCTACTACTGTCAACAGTATTATGAAA
AACCACATC

>rIGKV22U24 ; wgs
GACATCAAGATGACCCAGTCTCCTTCATTCCTGTCTGCATCTGTGGGAGACAGAGTCACTATCAACTGCA
AAGCAAGTCAGAATATTAACAAGTACTTAAACTGGTATCAGCAAAAGCTTGGAGAAGCTCCCAAACTCC
TGATATATAATACAAACAATTTGCAAACGGGCATCCCATCAAGGTTCAGTGGCAGTGGATCTGGTACTG
ATTTCACACTCACCATCAGCAGCCTGCAGCCTGAAGATGTTGCCACATATTTCTGCTTGCAGCATAATAGT
AGGCCG

>rIGKV22U25 ; wgs
GACATCCAGATGACCCAGTCTCCTTCATTCCTGTCTGCATCTGTGGGAGAAAGAGTCACTCTCAGCTGCA
GAGCAAGTCAGAATATTAACAAGTACTTAGACTGGTATCAGCAAAAGCTTGGAGAAGCTCCCAAACTCC
TGATATATAATACAAACAATTTGCATACGGGCATCCCATCAAGGTTCAGTGGCAGTGGATCTGGTACAG
ATTACACACTCACCATCAGCAGCCTGCAGCCTGAAGATGTTGCCACATATTTCTGCTTGCAGCGTAATAG
TTGGCCG

>rIGKV22U26 ; wgs
GACATCCAGATGACCCAGTCTCCTTCAGTCCTGTCTGCATCTGTGGGAGACAGAGTCACTCTCAACTGCA
AAGCAAGTCAGAATATTAACAAGTACTTAAACTGGTATCAGCAAAAGCTTGGAGAAGCTCCCAAACTCC
TGATATATAATACAAACAATTTGCAAACGGGCATCCCATCAAGGTTCAGTGGCAGTGGATCTGGTACAG
ATTACACACTCACCATCAGCAGCCTGCAGCCTGAAGATGTTGCCACATATTTCTGCTTTCAGCATAATAGT
TGGCCC

>rIGKV22U27 ; wgs
GACATCCAGATGACCCAGTCTCCTTCACTCCTGTCTGCATCTGTGGGAGACAGAGTCACTATCAACTGCA
AAGCAAGTCAGAATATTAACAAGTACTTAAACTGGTATCAGCAAAAGCTTGGAGAAGCTCCCAAACTCC
TGATATATAATACAAACAATTTGCAAACGGGCATCCCATCAAGGTTCAGTGGCAGTGGATCTGGTACAG
ATTACACACTCACCATCAGCAGCCTGCAGCCTGAAGATTTTGGCACATATTTCTGCTTTCAGCATAATAGT
TGGCCC

>rIGKV22U33 ; wgs
GACATCCAGATGACCCAGTCTCCTTCATTCCTGTCTGCAACTGTGGGAGACAGAGTCACTATCAACTGCA
AAGCAAGTCAGAATATTAACAAGTACTTAAACTGGTATCAGCAAAAGCTTGGAGAAGCTCCCAAACGCC
TGATATATAATACAAACAGTTTGCAAACTGGCATTCCATCAAGGTTCAGTGGCAGTGGATCTGGTACAG

ATTACACACTCACCATCAGCAGCCTGCAGCCTGAAGATGTTGCTACATATTTCTGCTTGCAGCATAATAG
TGGGCCC

>rIGKV22U53 ; wgs
AACATCCAGCTGACCCAGTCTCCTTCACTCCTGTCTGCATCTGTGGGAGACAGAGTCACTCTTAGCTGCA
AAGGAAGTCAGAATATTAACAATTACTTAGCCTGGTACCAACAAAAGCTTGGAGAAGCTCCCAAACTCC
TGATATATAATACAAACAGTTTGCAAACGGGCATCCCATCAAGGTTCAGTGGCAGTGGATCTGGTACAG
ATTACACACTCACCATCAGCAGCCTGCAGCCTGAAGATGTTGCCACATATTTCTGCTATCAGTATAACAA
CGGGTAC

>rIGKV22U72 ; wgs
GACATCCAGATGACCCAGTCTCCTTCACTCCTGTCAGCATCTGTGGGAGACAGAGTCACTCTCAGCTGCA
AAGCAAGTCAGAGTATTTACAGCAGCTTAGCCTGGTATCAGCAAAAGCTTGGAGAAGCTCCCAAACTCC
TGATATATAGTGCAAACAGTTTGCAAACGGGCATCCCGTCAAGGTTCAGTGGCAGTGGATATGGTACAG
ATTTCACACTCACCATCAGCAGCCTGCAGCCTGAAGATGTTGCCACATATTTCTGCCATCAGTATTACAGT
TGGCCC

>rIGKVxU54 ; wgs
GATATTCAAGTGACTCAATCTCCATCCTCCCTCTTGGCATCTCTAGGAGAGAGAGTCACTATCACATGCC
AGACAAGTCAGAGCATTAGCAATAACCTAAACTGGTATCAGCAGAAACCAGGACAAGCTCCTATGCTCT
TGATCTATTATGCAACCAGTTTGCAGACTGGCATGCCATCAAGGTTCAGTGGCCAATATTCTGGGAGAA
GTTTCACTCTAACCATCACTAGCCTGGAGCCAGAAGATATTGCAAATTATTTTTGTCTGCAGCATTACAGT
GCTCCTCA

>rIGKVxU84 ; wgs
GATATCCAGATGAACCAGACTCCATCTACTCTATCTATGTCTATTGGAGAACGAGTAATTATTAATTGCCA
TGCCAGTGAGATCATTAATACTTGGTTATCCTGGCACCAGCAGAAACCAGAGAATGCGCCACAACTACT
CATCTCTAAGGCATCCAACTTCCATACTGAGGTTCCATCAAGGTTCAGTGGCAGTGGATCTGGAACAGAT
TACTCTCTCAGCAGCAGCAGCCTCGAGCCTGAAGACATTGCTACTTACTACTGTGTACAGGCTAAGAGTC
TTCCTCC
